# Identifying spatially variable genes by projecting to morphologically relevant curves

**DOI:** 10.1101/2024.11.21.624653

**Authors:** Phillip B. Nicol, Rong Ma, Rosalind J. Xu, Jeffrey R. Moffitt, Rafael A. Irizarry

## Abstract

Spatial transcriptomics enables high-resolution gene expression measurements while preserving the two-dimensional spatial organization of the biological sample. A common objective in spatial transcriptomics data analysis is to identify spatially variable genes within predefined cell types or regions within the tissue. However, these regions are often implicitly one-dimensional, making standard two-dimensional coordinate-based methods less effective as they overlook the underlying tissue organization. Here we introduce a methodology grounded in spectral graph theory to elucidate a one-dimensional curve that effectively approximates the spatial coordinates of the examined sample. This curve is then used to establish a new coordinate system that reflects tissue morphology. We then develop a generalized additive model (GAM) to estimate spatial patterns which permits the detection of genes with variable expression in the new *morphologically relevant* coordinate system. Our approach directly models gene counts, thereby eliminating the need for normalization or transformations to satisfy normality assumptions. A second important advantage over existing hypothesis-testing approaches is that our method not only improves performance but also accurately estimates gene expression patterns and precisely pinpoints spatial loci where deviations from constant expression occur. We validate our approach through extensive simulation and by analyzing experimental data from multiple platforms such as Slide-seq and MERFISH. As an example of its ability to enable biological discovery, we demonstrate how our methodology enables the identification of novel interferon-related subpopulations in the mouse mucosa, as well as markers of inflammation-associated fibroblasts in a multi-sample spatial transcriptomic dataset.

## 1 Introduction

Spatial transcriptomics (ST) technologies permit high-resolution measurement of gene expression while maintaining the spatial coordinates of the samples (Rao et al., 2021; Moses and Pachter, 2022). These technologies have the potential to improve our understanding of the influence of cellular spatial organization on important biological processes and disease. One of the starting points for ST analysis is the identification of spatially variable genes (SVGs) (Adhikari et al., 2024). Since spatial variability can often be explained by differences in cell type (Cable et al., 2022b), it is common to test for SVGs within a predefined cell type or spatial domain (Yu and Luo, 2022).

Current statistical approaches for SVG detection perform a hypothesis test for each gene, quantifying the evidence of spatial variability using a *p*-value (Svensson et al., 2018; Sun et al., 2020; Hao et al., 2021; Zhu et al., 2021; Weber et al., 2023). However, these methodologies are incapable of distinguishing genes whose spatial expression patterns manifest in fundamentally different ways, such as along distinct anatomical features within the tissue. For example, MERFISH measurements of a healthy mouse colon (Xu et al., 2026) revealed two dominant patterns of spatial variability, denoted here as *localized* (**Fig 1a, left**) and *radial gradient* (**Fig 1a, right**), respectively. Genes exhibiting localized variation, such as *Ddx58*, are characterized by a distinct patch of expression in one region of the colon, whereas genes exhibiting radial gradient variation, such as *Apob*, are characterized by a gradual change in expression between the outside and inside of the mucosa. Importantly, while both of these examples are illustrations of spatially variable genes, their distinct spatial distributions have important implications for their biological interpretation. *Ddx58* is an interferon-stimulated gene, and the localized expression observed in the mucosa is representative of local patches of interferon activity and interferon-stimulated gene expression described previously (Van Winkle et al., 2022). By contrast, the radial distribution of *Apob*, a marker of the final stages of mature enterocytes, shows the known variation of epithelial cells from the base to the tip of colonic crypts (Moor et al., 2018). Although current leading approaches successfully identified these genes as spatially variable (**Fig S1, S2**), they lack the capability of distinguishing between these two modes of spatial variation. Additionally, these existing methodologies do not allow for precise pinpointing of the locations where spatially relevant gene expression occurs.

**Figure 1.**
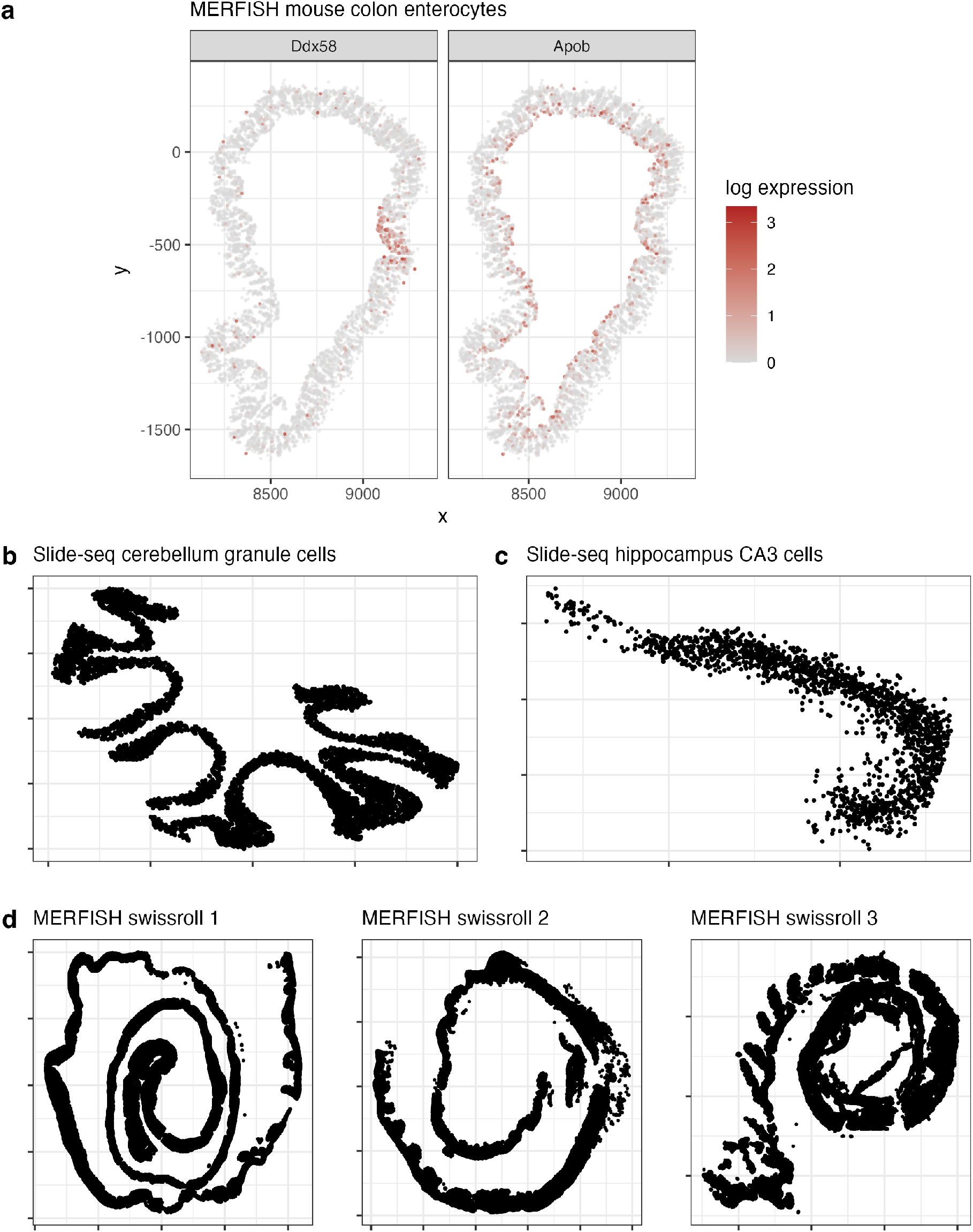
**a.** The spatial location of enterocytes identified in a cross section of the healthy mouse colon as measured with MERFISH. Cells are colored by the log-transformed expression of the two listed genes. The expression of *Ddx58* is called localized whereas the expression pattern of *Apob* is called radial gradient. The spatial locations of the plotted enterocytes lie close to a one-dimensional circular manifold. Additional examples of cell types with coordinates that lie close to a one-dimensional curve can be found in **b**. granule cells from the mouse cerebellum (Cable et al., 2022b) and **c**. CA3 cells in the mouse hippocampus (Stickels et al., 2021). **d**. Three mouse colon swiss rolls from the MERFISH study of Cadinu et al. (2024), details in Methods.

Although numerous statistical techniques exist for the estimation of two-dimensional surfaces (Wood, 2003; Schulz et al., 2018), and some of these are used for ST (Cable et al., 2022a), it is noteworthy that in many applications the primary interest is in genes that exhibit variation along one-dimensional paths. For example, the exercise of visually detecting the localized pattern shown in *Ddx58* could be described as searching for deviation from a baseline expression level along a curve tracing through the mucosa. The radial gradient pattern exhibited by *Apob* could be described as change in expression in the direction perpendicular to the curve. This implies that a curve-based coordinate transform could help separate genes with a localized burst from those with radial gradient, which, in turn, facilitates new biological findings. In addition to the colon (**Fig 1a**), cell types in the brain also commonly exhibit distinct one-dimensional spatial structure. We therefore also consider two Slide-seq datasets, granule cells from the mouse cerebellum (Cable et al., 2022b) (**Fig 1b**) and CA3 cells from the mouse hippocampus (Stickels et al., 2021) (**Fig 1c**), and a MERFISH study consisting of three mouse colons organized in a Swiss roll structure (Cadinu et al., 2024) (**Fig 1d**).

We introduce a statistical framework that estimates a one-dimensional curve passing through the ST coordinates and then uses that estimated curve to define a *morphologically relevant* coordinate system. Although similar curve-estimation methods have been used for pseudotime analysis in single-cell RNA-seq (Street et al., 2018), we find that our methodology grounded in spectral graph theory yields better results on two-dimensional ST data. Moreover, pseudotime methods do not measure variation *orthogonal* to the curve which is critical to distinguish, for example, between localized and radial gradient patterns. We also compare this approach to several existing ST methodologies that leverage one-dimensional structure such as GASTON (Chitra et al., 2025), Spateo (Qiu et al., 2024) and computer vision techniques such as skeletonization (Maragos and Schafer, 2003), finding that our explicit model of morphology leads to better estimation and downstream results.

Upon estimating the curve, we pose a generalized additive model (GAM) to model expression as a (possibly non-linear) function of the morphologically relevant coordinates. We refer to our approach as *MorphoGAM*. A key strength of MorphoGAM is that it estimates an entire spatial expression function, thereby allowing the precise localization of spatially interesting expression. We also show that, in general,

MorphoGAM increases the statistical power to detect scientifically relevant SVGs. This increased power translates to improved biological findings on the mouse mucosa data by demonstrating that MorphoGAM identifies interferon-related gene sets that are missed by existing SVG methods. Finally, we apply MorphoGAM to the MERFISH swiss rolls, and identify novel markers of inflammation-associated fibroblasts.

## 2 Results

### MorphoGAM identifies morphologically relevant coordinates in spatial transcriptomics data

We begin by modeling the spatial location of cell *j*, 1 ≤ *j* ≤ *n*, using *morphologically relevant* coordinates *t*_*j*_ and *r*_*j*_. Specifically, we assume that the standard two-dimensional spatial coordinates *x*_*j*_ ∈ ℝ^2^ lie close to a latent curve:

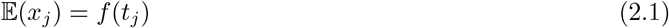

where *f* : [*a, b*] → ℝ^2^ is a smooth one-dimensional parametric curve. We write *f* (*t*) = (*f*_1_(*t*), *f*_2_(*t*)) to denote the two component functions of the curve. The first morphologically relevant coordinate *t*_*j*_ describes the position of cell *j* along the curve. In the Methods we describe in detail our approach based on spectral graph theory to estimate *t*_*j*_ and *f* . Briefly, our approach relates the distance between coordinates |*t*_*i*_ − *t*_*j*_| to the shortest path in a *k*-nearest neighbor graph *G*_*k*_ and then shows that *t*_*j*_ can be estimated through an eigendecomposition of the centered shortest path matrix. After obtaining an estimate 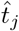, the curve *f* can be estimated by smoothing each dimension separately. We plug in 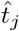 to (2.1) to obtain

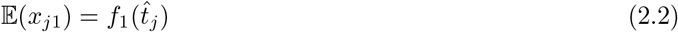

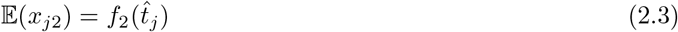

We thus obtain 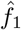 and 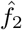 by using regression splines as implemented by Wood (2017). Although we find the estimated curve is typically robust to the choice of *k*, we provide in Methods a model selection procedure to automatically select *k*.

Our methodology to estimate *t*_*j*_ in the case of an open curve (*f* (*a*) ≠ *f* (*b*)), is motivated by the ISOMAP technique (Tenenbaum et al., 2000) for non-linear dimensionality reduction. We extend this approach to allow our method to address scenarios wherein *f* constitutes a closed curve (*f* (*a*) = *f* (*b*)). A detailed visual assessment indicates that, when applied to the slice of healthy mouse colon, MorphoGAM excels in estimating *f* (*t*) (**Fig 2a**) and the morphologically relevant coordinate *t*_*j*_ (**Fig 2b**).

**Figure 2.**
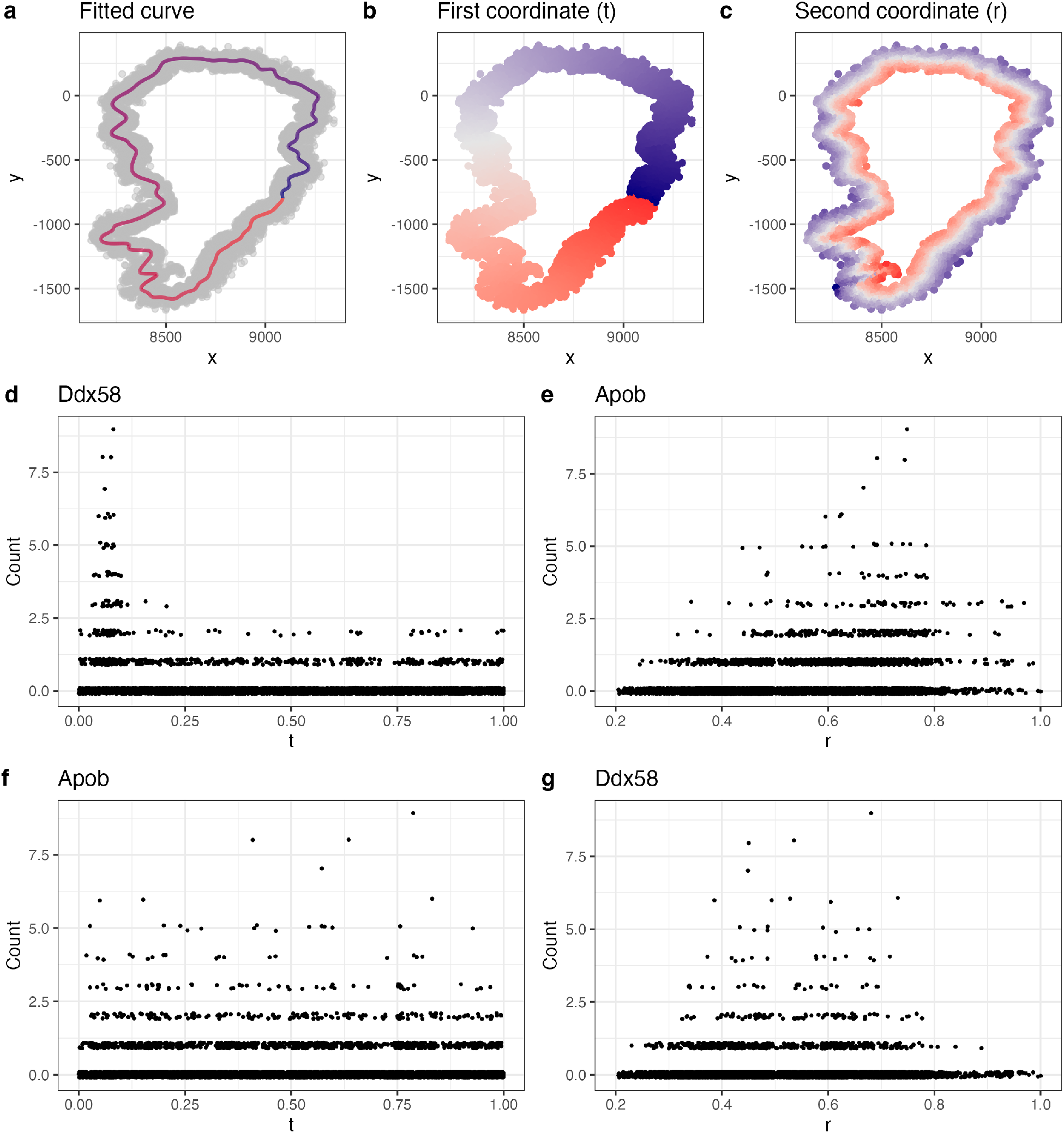
Overview of MorphoGAM. **a.** MorphoGAM begins by estimating a smooth parametric curve passing through the spatial transcriptomic sample coordinates. **b**. The first morphologically relevant coordinate is defined as the position of each cell (or more generally, sample) along the estimated curve from the previous step. **c**. The second morphologically relevant coordinate is defined as the position of the cell in the direction orthogonal to the curve at a given point. **d**. Genes with localized variation such as *Ddx58* show strong expression variation as a function of the first morphologically relevant coordinate. **e**. Genes with a radial gradient pattern such as *Apob* show expression variation as a function of the second morphologically relevant coordinate. **f**. and **g**. The same genes as a function of the other morphologically relevant coordinate do not show significant spatial variation.

The second *morphologically relevant* coordinate, denoted here by *r*_*j*_, is defined by using the distance from the cell’s coordinates to the position on the curve *f* (*t*). Explicitly, the magnitude of *r*_*j*_ is given by

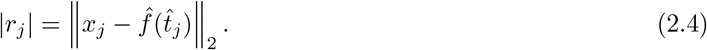

To determine the sign of *r*_*j*_, we set

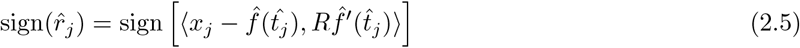

where *R* : ℝ^2^ → ℝ^2^ is a counter-clockwise 90 degrees rotation: *R*(*v*_1_, *v*_2_) = (−*v*_2_, *v*_1_). The conceptual framework behind equation (2.5) can be understood by envisioning a traversal along the curve, where the velocity vector at time *t* is *f*^*′*^(*t*). Residuals exhibiting a positive sign would be placed on the left-hand side of the curve as one progresses, whereas those with a negative sign would be on the right-hand side. The left-hand side can be ascertained through a counter-clockwise rotation *R* of the velocity vector *f*^*′*^(*t*). This coordinate for each cell is also morphologically pertinent, as illustrated in Figure 2c.

After transforming to the morphologically relevant coordinate system, the difference between localized and radial gradient patterns becomes immediately clear. SVGs with localized patterns show variation in the first morphologically relevant coordinate *t*_*j*_ (**Fig 2d**) whereas SVGs exhibiting a radial gradient pattern show variation in the second morphologically relevant coordinate *r*_*j*_ (**Fig 2e**). Importantly, the genes with strong localized or radial gradient pattern show little spatial variation with respect to the other morphologically relevant coordinate (**Fig 2f-g**).

### MorphoGAM outperforms existing curve estimation approaches

Hastie and Stuetzle (1989) introduced model (2.1) as a general approach to estimate a curve passing through a set of points (in arbitrary dimension). This method, known as *Principal Curves*, is used by the popular pseudotime method *Slingshot* (Street et al., 2018). Hastie and Stuetzle (1989) employ an iterative algorithm that alternates between updating *f* and updating *t*_*j*_. However, we find that this iterative approach is unsuitable for the highly non-linear structures observed in spatial transcriptomics data. To demonstrate this, we applied *Principal Curves* to granule cells from the mouse cerebellum, measured using Slide-seqV2 (Cable et al., 2022b) (see Figure 1b). We found that this approach did not accurately estimate the curve for a variety of tuning parameter choices (**Fig 3a-c**). In contrast, the automatically selected value of *k* = 9 accurately identified the curve and the morphologically relevant coordinate (**Fig 3d**). The performance was nearly identical for *k* = 5 (**Fig 3e**) and only began to degrade once *k* ≥ 30 (**Fig 3f**).

**Figure 3.**
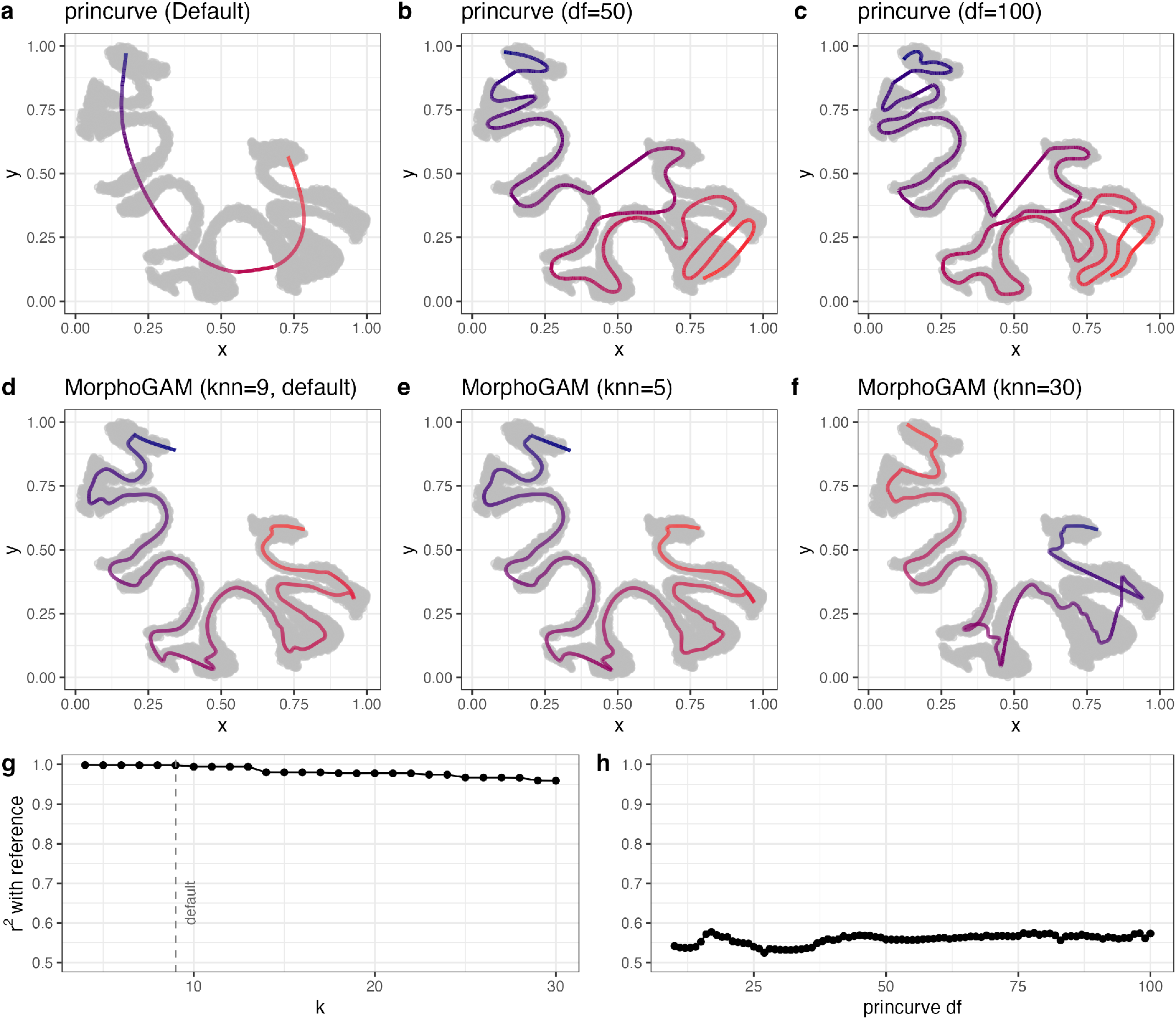
MorphoGAM outperforms existing curve estimation approaches. **a.** Applying the principal curves method (Hastie and Stuetzle, 1989) as implemented by the *princurve* (Cannoodt, 2018) R package. **a**., **b**., and **c**. show the estimated curve for three different values of the tuning parameter, which is degrees of freedom (df) of the smoothing spline. **d**., **e**. and **f**. show the estimated curve from MorphoGAM with three different values of its tuning parameter k-nearest neighbor (kNN). **g**., **h**. Using the estimated coordinate *t*_*j*_ from a hand-drawn reference curve (**Fig S3b**) we compute the squared Spearman correlation between this and the estimated coordinate from both methods as the tuning parameters vary.

To quantitatively evaluate the robustness of these methods with respect to their tuning parameters, we defined a *reference coordinate t*_*j*_ by carefully hand-drawing a path (**Fig S3**) and compared estimates 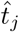 obtained with different values of the tuning parameter to the *t*_*j*_. We observed that the Spearman correlation between the estimates 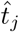 and *t*_*j*_ was below 0.6 for Principal Curves (**Fig 3g**). In contrast, with MorphoGAM, the correlation exceeded 0.95 for *k <* 10 and remained above 0.9 for values up to *k* = 30, demonstrating robustness across choices of *k* (**Fig 3h**). We also applied both methods to simulated Swiss roll data (**Fig S4**) and the mouse colon dataset (**Fig S6**) and found similar performance gains from using MorphoGAM. An additional advantage of MorphoGAM is that the tuning parameter *k* (number of nearest neighbors) is more interpretable than degrees of freedom (df) for a smoothing spline, which makes it easier to find a reasonable value in practice. To quantify the performance of the *r* coordinate estimation, we applied it to simulated annulus datasets and found strong performance across a range of tuning parameters (**Fig S5**).

Next, we compared MorphoGAM to existing ST methods that do not explicitly estimate a curve, such as GASTON (Chitra et al., 2025), a method that uses deep learning to estimate a one-dimensional value (termed *isodepth*) that summarizes tissue topography. Although the isodepth is noticeably higher near the region where *Ddx58* has an increase in expression, the isodepth does not perfectly correspond to either of the morphologically meaningful directions (**Fig S7**) and both patchy and gradient genes show variable expression as a function of the isodepth (**Fig S8**), showing that a single dimension is insufficient to separate the biologically distinct group of genes. We also compared MorphoGAM to the spatial digitization method Spateo (Qiu et al., 2024), which estimates a coordinate by solving Laplace’s equation between two manually selected poles. Although this approach has been demonstrated on brain data similar to the granule dataset, we found that it did not adequately capture the loop structure of the mucosa, assigning distant points the same underlying coordinate (**Fig S9**).

Finally, we compared the performance of MorphoGAM to a morphological skeleton, a computer vision technique used to find thin structures in images (Maragos and Schafer, 2003). Briefly, the ST coordinates from the granule cells were converted to a 512 × 512 binary image, and *magick* (Ooms, 2025) was applied to identify the skeleton. Although the skeleton generally captures the 1D structure of the coordinates, it does not explicitly fit a latent curve and thus does not permit the definition of a morphologically relevant coordinate system (**Fig S10**).

### MorphoGAM allows for interpretable detection of spatially variable genes

Following the estimation of morphologically relevant coordinates 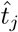 and 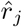, MorphoGAM identifies spatially variable genes using a generalized additive model (GAM) (Hastie and Tibshirani, 1986). Specifically, we denote the count for gene *g* in cell *j* with *Y*_*gj*_ and model it with

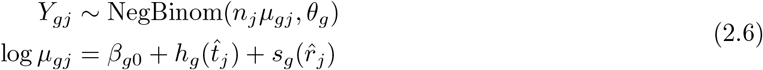

where *h*_*g*_ and *s*_*g*_ are unknown smooth functions, *β*_*g*0_ is an unknown intercept, *n*_*j*_ is the total counts for cell *j*, and *θ*_*g*_ is the inverse dispersion parameter.

In this model, gene *g* is spatially variable if *h*_*g*_≠ 0 or *s*_*g*_≠ 0. Estimating the parameters of model (2.6) is achieved by writing *h*_*g*_ and *s*_*g*_ as the sum of basis functions and then adding a penalty to encourage smoothness (Methods). Although *p*-values can be computed by testing the null hypothesis *h*_*g*_ = 0 or *s*_*g*_ = 0, we recommend inspecting the estimated functions *ĥ*_*g*_ and *ŝ*_*g*_ along with estimates of their covariance to measure uncertainty. To avoid misinterpreting noisy estimates as true biological signal, we opt to shrink (pointwise) each function towards 0 by an amount related to its standard error (see Methods).

Although users may wish to examine the entire function estimate, there is often a summary (or *functional*) that could be more useful for addressing the particular scientific question. For example, the *peak* summary could be used to quickly identify genes that differ significantly from baseline (on the log-fold change scale):

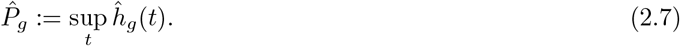

which estimates the maximum log-fold change from the baseline log expression 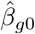. Because this measurement could prioritize large multiplicative changes in small genes we also define the *range*

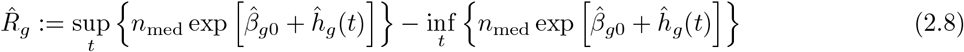

which can identify genes that have large differences on the original scale of the counts. Here *n*_med_ is defined as the median of the *n*_*j*_, so that 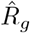 can be directly interpreted as a count difference. We note *ŝ*_*g*_(*t*) could replace *ĥ*_*g*_ (*t*) in both equations (2.7) and (2.8).

We emphasize that model (2.6) can easily be modified depending on the particular scientific question. For example, if only variation along the curve is of interest then we only need to examine *ĥ*_*g*_ (*t*). Moreover, the model is flexible enough to account for other potential confounders in the linear predictor.

### MorphoGAM improves power to detect relevant spatially variable genes

We applied MorphoGAM to CA3 cells from the mouse hippocampus (**Fig 1c**) to estimate a one-dimensional curve 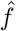 and morphologically relevant coordinates 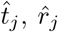 (**Fig 4a**). In the original analysis of this dataset, Cable et al. (2022b) used 2D locally weighted regression to identify genes with a high coefficient of variation (CV). This analysis identified two genes *Rgs14* and *Cpne9* that exhibited variable expression near opposite ends of the hippocampal structure. We then applied the GAM model (2.6) to identify genes varying along the curve 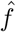. Our approach corroborated the finding of *Rgs14* (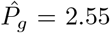, *p <* 10^−6^) and *Cpne9* (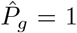, *p <* 10^−6^) (**Fig 4b**). When ranking all genes by the peak or range statistic, our approach identified genes that were not reported in the original analysis of Cable et al. (2022b) such as *Fxyd6* (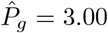, *p <* 10^−6^) and *Hpca* (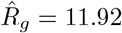, *p <* 10^−6^) (**Fig 4c**).

**Figure 4.**
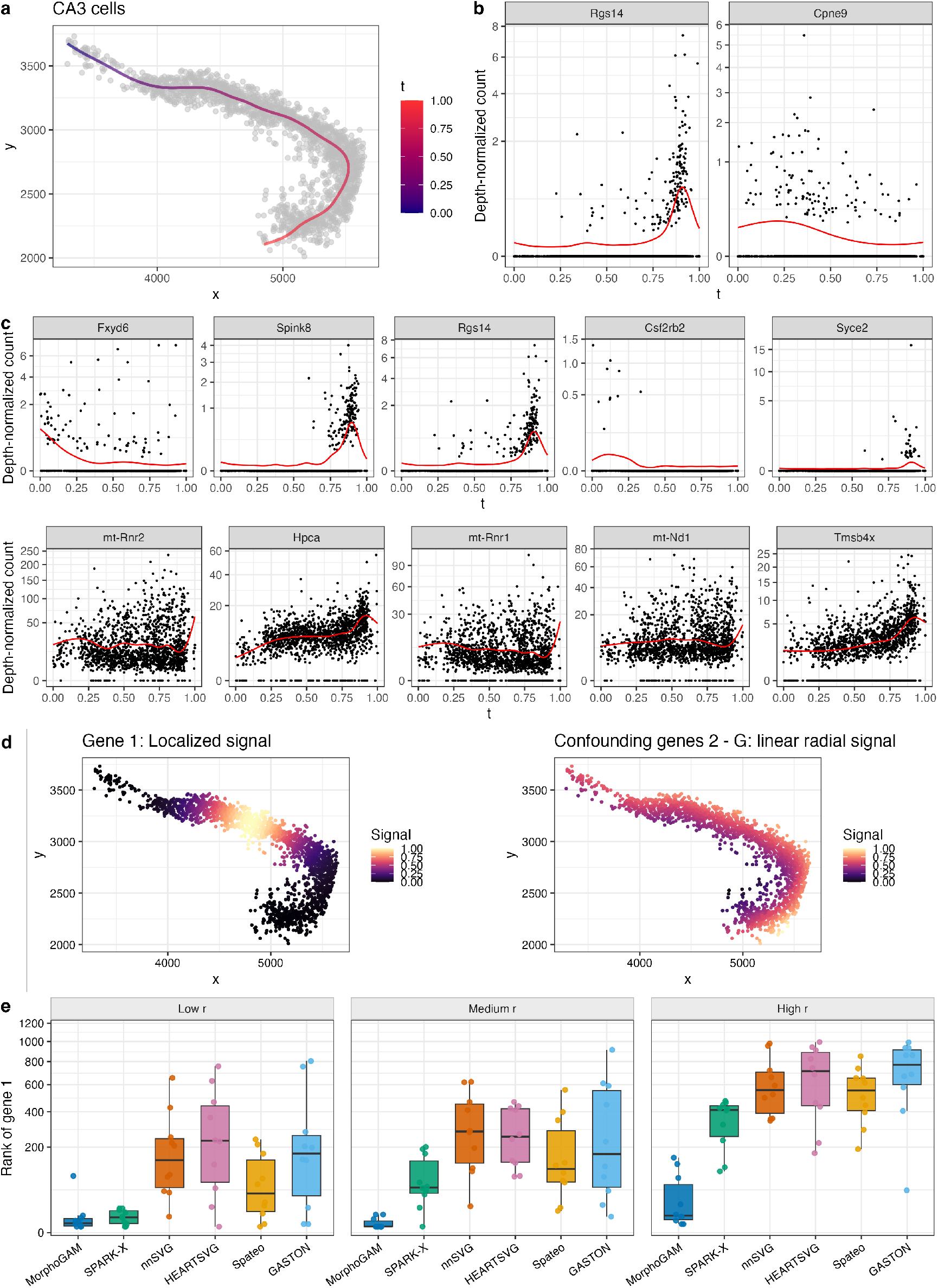
MorphoGAM increases power to detect relevant spatially variable genes. **a.** The estimated curve 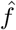 on the CA3 cells from mouse hippocampus (see **Fig 1c**). **b**. The estimated functions 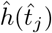 for two previously reported spatially variable genes *Rgs14* and *Cpne9*. **c**. Plotting the genes with the largest peak and range summaries. **d**. Our simulation study considered a single gene with variation in the *t* direction as well as many genes with variation in the *r* direction. **e**. The ranking of gene 1 (see Methods) for each of the methods, as the variation in the *r* direction is increased.

Starting from the spatial coordinates of the CA3 cells, we conducted a broad simulation study to demonstrate the increased power to identify genes with variable expression along the morphological trajectory. Specifically, we generated counts from a single gene with variable expression along the *t* direction as well as *G* − 1 genes with variable expression along the *r* direction (**Fig 4d**, Methods). Supposing the scientific goal involves identifying variation in the *t* direction (variation along the curved hippocampal structure), we assess the performance of MorphoGAM and five other methods to recover the *t*-associated gene. As each method can produce a ranked list of candidate SVGs (Methods), we use the rank of the *t*-associated gene as a metric for MorphoGAM’s performance. We considered 3 scenarios (low *r*, medium *r*, high *r*) pertaining to the signal strength of the *r* genes, reflecting our motivating example (**Fig 1a**) where the signal of the patchy genes was washed away by the numerous radial gradient genes. The results in Figure 4e demonstrate that, although the other methods can perform well in the low *r* scenario, performance degrades in the high *r* scenario. On the other hand, MorphoGAM decouples *t* and *r* variation and performs strongly in all scenarios. Figures S11 and S12 repeat the simulation with different expression patterns for the *t* gene as well as different spatial coordinates to demonstrate MorphoGAM’s flexibility to capture arbitrary patterns of spatial expression across different morphologies. In Supplementary section S2.1, we also describe and perform a more traditional statistical power and type I error analysis.

### MorphoGAM identifies groups of spatially co-varying genes in the mouse colon data

We applied MorphoGAM to the MERFISH measurements of a slice of healthy mouse colon (**Fig 1a**) with the goal of separating genes with localized and radial gradient patterns of expression. Because localized genes are characterized by a burst in expression along the curve, we used the peak of estimated functions 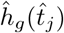 to rank genes (**Fig 5a**). Radial gradient genes, on the other hand, are characterized by a smooth transition along the second morphologically relevant coordinate, so for this we found genes with a large range in *ŝ*_*g*_ (**Fig 5b**). **Figure 5** also lists the ranking of each gene of nnSVG, SPARK-X, GASTON, and Spateo, showing that the targeted analysis of MorphoGAM prioritizes genes that could have been missed if alternative methods were used to identify SVGs. In particular, *Ddx58* was found to have a large peak in the first morphologically relevant coordinate (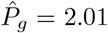, *p <* 10^−6^) and *Apob* was found to have a large peak in the direction of the second morphologically relevant coordinate (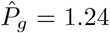, *p <* 10^−6^). We also plot the genes with the largest range in the direction of the first morphologically relevant coordinate and the genes with the largest peak in the direction of the second morphologically relevant coordinate in **Figure S13**. We repeated the analysis using curves with different values of *k* as well as a manually drawn curve and found that the resulting SVG rankings were highly consistent, indicating that the analysis is robust to the precise specification of the curve *f* (**Fig S14)**. To quantitatively compare the list of genes identified by the different SVG algorithms, we performed gene set analysis (GSA) (Subramanian et al., 2005; Väremo et al., 2013) using the mouse hallmark gene sets (Liberzon et al., 2015; Castanza et al., 2023). For MorphoGAM, we ranked genes by their peak statistic in the *t*-coordinate and found that, as expected, the most enriched gene sets corresponded to interferon activity (**Fig 5c**). On the other hand, the enriched gene sets from the other methods included other gene sets associated with radial genes, showing that the focused analysis of MorphoGAM can lead to improved biological findings (**Fig 5c**).

**Figure 5.**
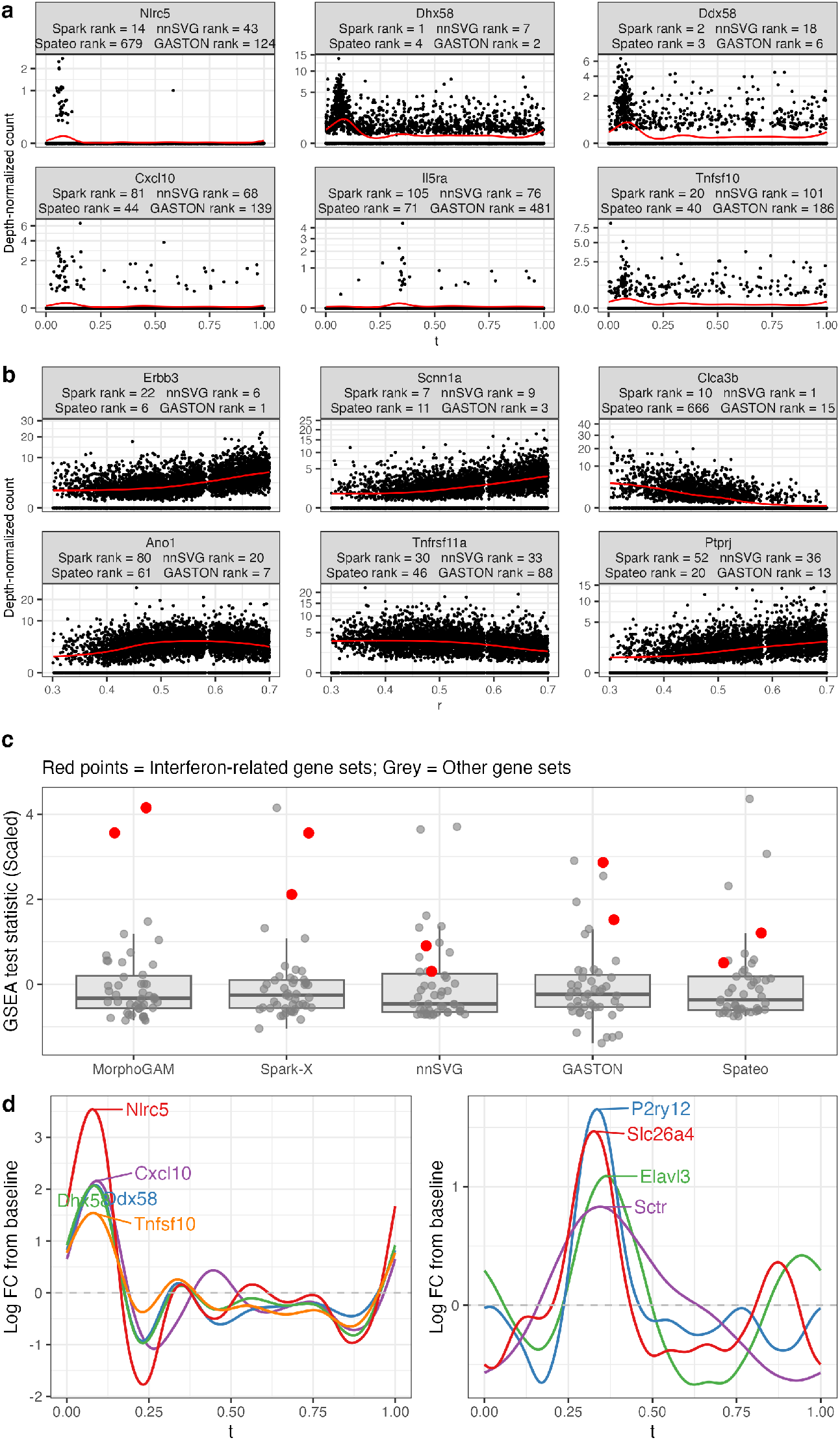
MorphoGAM identifies additional genes with localized and radial gradient patterns. **a.** The top six genes identified when ranking by the peak of *h*_*g*_. That is, genes with a high log fold-change relative to baseline in the direction of the first morphologically relevant coordinate. **b**. The top six genes identified when ranking by the range of *s*_*g*_. That is, genes with a large change on the scale of the counts in the direction of the second morphologically relevant coordinate. Each label shows the ranking of the gene from SPARK-X, nnSVG, Spateo, and GASTON. **c**. We used the piano R package (Väremo et al., 2013) to perform gene set analysis using mouse hallmark gene sets. For each method, we scaled the GSA test statistics to have mean zero and unit variance and highlighted the interferon-related gene sets in red. **d**. We used singular value decomposition (see Methods) to identify groups of genes with similar spatial patterns. The top two groups of genes are plotted together as a function of *t*.

Because MorphoGAM provides quantitative estimates of gene expression spatial patterns, pattern discovery techniques, such as singular value decomposition (SVD), can be readily applied to identify groups of genes with spatially similar expression patterns along the one-dimensional path (see Methods). The first group identified the patch interferon-stimulated genes *Cxcl10, Ddx58, Dhx58, Tnfsf10* (Van Winkle et al., 2022), as well as *Nlrc5*, a negative regulator of the interferon pathway (Cui et al., 2010) (**Fig 5d**, left). The second group identifies neuro-related genes *P2ry12* (Gómez Morillas et al., 2021), *Slc26a4, Ptpro* (Jiang et al., 2017), *Elavl3* (Scheckel et al., 2016), suggesting a local region of neuronal infiltration at *t* ≈ 0.3 (**Fig 5d**, right).

### MorphoGAM identifies markers of inflammation-associated fibroblasts

Because MorphoGAM is based on a statistical model, it also facilitates the integration and analysis of multi-sample spatial transcriptomic data sets with shared spatial structures. To demonstrate, we considered a MERFISH dataset consisting of three *Swiss rolls* from a mouse colitis model (Cadinu et al., 2024), (**Fig 1d**). We found that the overlapping points led to a *k*-NN graph that could not accurately capture geodesic distances; as a result, we opted to hand-draw the curve *f* (*t*) using the interactive tool described in Methods (**Fig 6a**). To identify genes associated with inflammation-associated fibroblasts (IAFs) in the colon we focused on ulcerated regions that span the longitudinal direction of the colon (**Fig 6b**), and fit the following modification of model (2.6):

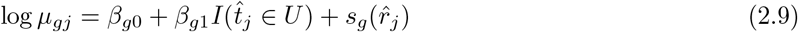

where the indicator function *I*(*t*_*j*_ ∈ *U*) = 1 if cell *j* is in the ulcerated region and is 0 otherwise. We used this model to identify genes with large effect size *β*_*g*1_ across all three Swiss rolls, indicating that the gene is a persistent marker of an IAF (see S2.2). This analysis identified IAF markers previously identified by (Cadinu et al., 2024) such as *Grem1, Il11, Mmp3*, and *Mmp10* (**Fig 6c**). MorphoGAM also identified *Timp1*, which was not identified in the original analysis of these data, but is known to play a role in inflammation (Wang et al., 2022), highlighting the increased sensitivity of MorphoGAM and its ability to extend to multi-sample spatial transcriptomic studies.

**Figure 6.**
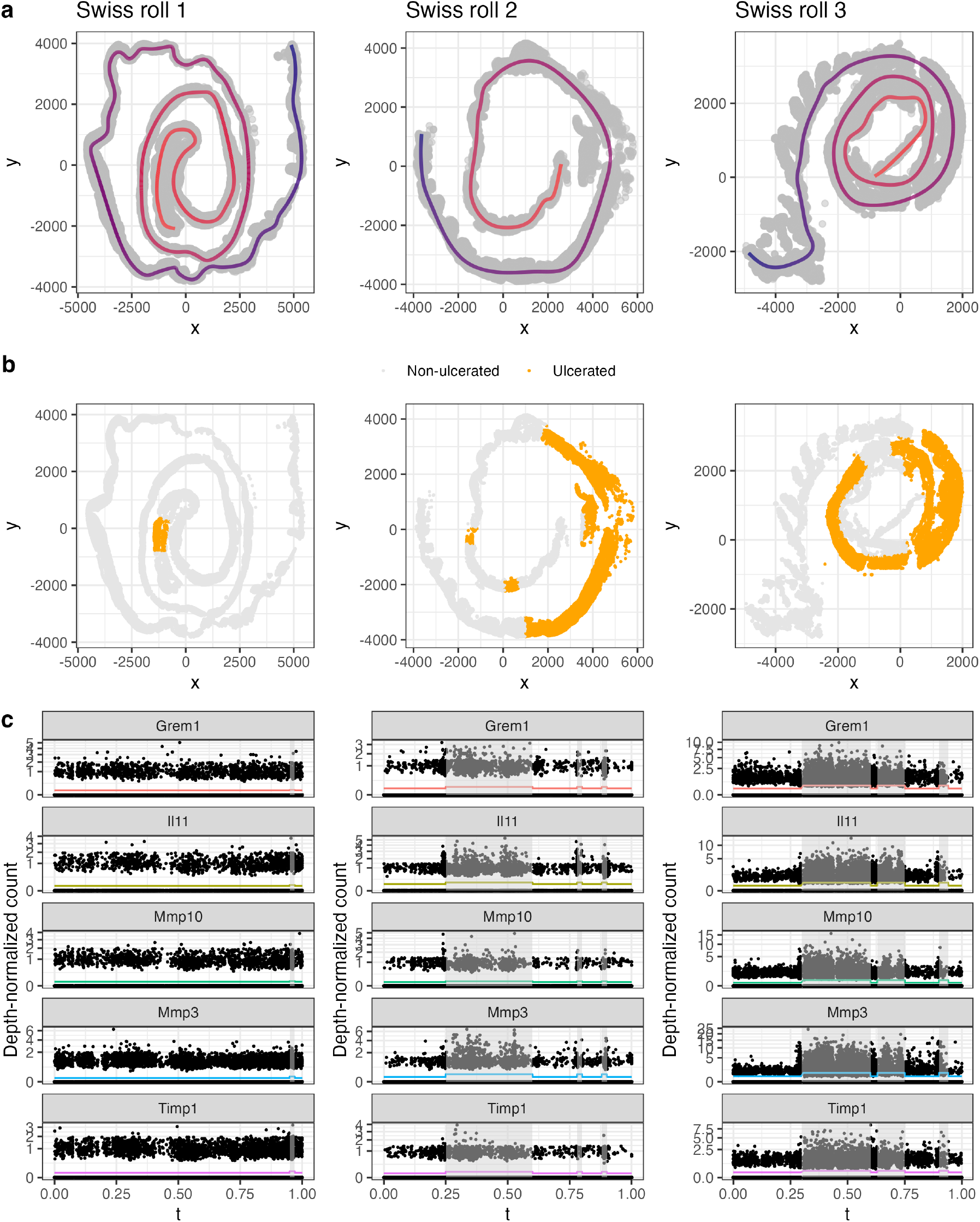
**a.** Three MERFISH colon swiss rolls from a mouse colitis model obtained from (Cadinu et al., 2024). The MorphoGAM R package was used to interactively draw the curve *f* (*t*) parameterizing the trajectory of the swiss rolls. **b**. The ulcerated regions in each swiss roll (highlighted in gold) manually identified by comparing to Figure S4 of (Cadinu et al., 2024) and by inspecting regions where *Vil1* expression is lower or absent. **c**. Markers of IAF were identified using the procedure in S2.2 and plotted as a function of *t*. The vertical shading indicates ulcerated regions and the colored line plots *n*_med_ exp(*β*_*g*0_ + *β*_*g*1_*I*(*t*_*j*_ ∈ *U*)).

### MorphoGAM can be used with any spatial transcriptomic technology

The statistical formulation of MorphoGAM is general and can be applied to any ST technology that provides spatial coordinates *x*_*i*_ with corresponding gene counts *y*. Beyond the Slide-seq and MERFISH datasets analyzed above, we evaluated MorphoGAM on two additional widely used ST platforms: 10X Genomics Xenium and Visium.

Applied to Xenium data from the mouse hippocampus (**Fig. S15a**), MorphoGAM captured spatial variation across CA1, CA2, and CA3 neurons. Curve estimation was used to model the intrinsic one-dimensional structure, and the previously described SVD approach clustered genes with shared spatial patterns. The first gene module delineated CA1 and CA3 regions (**Fig. S15b**). MorphoGAM identified markers such as *Fibcd1* and *Satb2*, which were not reported in a comparable analysis (Fu et al., 2024) but are validated CA1 markers in the Allen Brain Atlas (Lein et al., 2007).

We next analyzed a Visium dataset of the dorsolateral prefrontal cortex (Maynard et al., 2021), which includes manual annotations of cortical layers 1–6 (**Fig. S16a**). MorphoGAM identified a curve within layer 3 and projected the remaining spots onto this axis, yielding a continuous coordinate system spanning all layers (**Fig. S16b**). We applied a count-based extension of PCA (scGBM, Nicol and Miller (2025)) to these data and observed that the top two principal components varied smoothly as a function of the MorphoGAM *r* coordinate (**Fig S16c**). This suggests that gene expression varies continuously with cortical depth rather than strictly following laminar boundaries. By identifying genes with significant variation in the *r* direction, MorphoGAM recovered established layer-specific markers, including *AQP4, TRABD2A, HPCAL1*, and *KRT17* (all *p <* 10^−16^, all peak statistics *>* 0.8) (**Fig. S16d**), and further identified novel SVGs such as *GFAP* (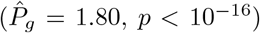, *p <* 10^−16^), a known astrocytic marker (Hol and Pekny, 2015) showing focal upregulation in a thin subset of layer 6 (**Fig. S16e**). These findings highlight the utility of the MorphoGAM *r* coordinate as a continuous descriptor of cortical depth, complementing or replacing discrete laminar annotation.

### MorphoGAM can be applied outside one-dimensional geometries

MorphoGAM is primarily designed for situations where the tissue coordinates are well approximated by a one-dimensional manifold and is not expected to be immediately applicable to general two-dimensional layouts. We note, however, that MorphoGAM can be applied beyond strictly one-dimensional manifolds when a biologically motivated direction of interest can be defined. To demonstrate, we apply MorphoGAM to a Visium dataset of a solid tumor (melanoma) from a zebrafish sample (Hunter et al., 2021). Although a solid tumor is not a one-dimensional structure, MorphoGAM (with a hand-drawn curve) tracing the boundary of the tumor leads to a new coordinate system where *t* reflects the position along the tumor boundary and *r* the position along the boundary-to-core axis (**Fig S17a, b**). We then apply the GAM model to identify collections of genes that vary with respect to both coordinates (see S2.3 for additional details). In the *t* direction, we identified a collective burst of many genes where the tumor contacts the surrounding muscle cells (**Fig S17c**). Functional enrichment analysis (Kolberg et al., 2020) of the top genes in this factor revealed over-representation of genes associated with degradation of the extracellular matrix, consistent with tumor microenvironment remodeling at the interface (**Fig S17d**). On the other hand, identification of genes with variation in the *r* direction yielded *hsp70l* (**Fig S17e**,**f**), a heat shock protein whose expression can reflect hypoxia or other forms of cellular stress in the tumor core (Sherman and Gabai, 2015).

## 3 Discussion

We introduced an approach to estimate the curve passing through spatial transcriptomics coordinates and leveraged this curve to define *morphologically relevant* coordinates. A model-based approach was used to quantify spatial variation along these morphologically relevant coordinates, which we have shown to be an interpretable and powerful approach to find relevant spatially variable genes. Importantly, we have advocated to directly use summaries of the estimated functions rather than relying on a null hypothesis test, as *p*-values do not provide information about the mode of spatial variation and are in general sensitive to misspecification in the assumed model (Greenland et al., 2016). MorphoGAM also permits the discovery of groups of genes with spatially correlated patterns, uncovering spatially co-localized cell subpopulations without the need for clustering techniques. Finally, we demonstrated that MorphoGAM can integrate and analyze multiple spatial transcriptomic samples when each sample has similar morphology.

While MorphoGAM currently relies on accurate annotation of cell types or spatial domains, this dependency highlights an exciting opportunity to integrate improved cell-type annotation tools and spatial domain discovery methods in future work. Similarly, although our approach is optimized for tissues that can be represented along a one-dimensional framework, this opens the door to flexible user-driven extensions. For example the accompanying software includes a manual curve-drawing tool that enables MorphoGAM to be applied even in cases where predetermined trajectories are of interest (as was demonstrated in Figure S17). We note, however, that the current implementation of MorphoGAM is most suitable when the latent curve is non-intersecting (except possibly at the endpoints). Self-intersecting or branching geometries (such as tissue organized around bifurcating ducts or blood vessels) introduce an ambiguity because locations near an intersection do not have a unique coordinate along a single curve. Future work could extend the automatic estimation procedure to branching structures or other violations of the one-dimensional manifold assumption. In summary, the underlying GAM framework is naturally extensible, and the same principles, summarizing estimated functions by range, peak, and other features—can be adapted to two-dimensional domains using thin-plate splines (Wood, 2003), broadening the scope of applications beyond the current setting.

Finally, we note that morphologically relevant coordinates may offer considerable utility in multi-sample ST analyses. For example, our analysis on the MERFISH swiss rolls (**Fig 6**) shows that, in settings where the samples have a shared morphology and identifiable endpoints, MorphoGAM can be used to compare samples using their *t* coordinate without first aligning their two-dimensional spatial coordinates. Future research could systematically extend the application of morphologically relevant coordinates to conduct multi-sample ST analyses.

## 4 Methods

### Statistical model for latent curve

Let *x*_*j*_ ∈ ℝ^2^ denote the spatial coordinates of cell 1 ≤ *j* ≤ *n*. We assume that *x*_*j*_ is a random variable with mean satisfying the following equation:

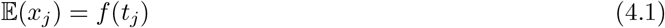

where *f* : [*a, b*] → ℝ^2^ is an (unknown) smooth parametric curve. We will write *f* (*t*) = (*f*_1_(*t*), *f*_2_(*t*)) to denote the two component functions of the curve. For the moment, we assume that *f* does not intersect itself, so that *t*_*i*_≠ *t*_*j*_ implies *f* (*t*_*i*_) *f* (*t*_*j*_). Note that both *f* and *t*_*j*_ are unknown and must be estimated. See Section S1.2 for detailed discussions on the identifiability conditions for *f* and *t*_*j*_.

The arclength of *f* can be expressed in terms of the first derivative 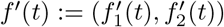:

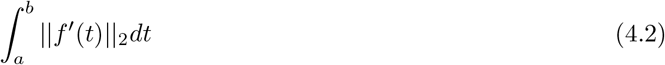

We will assume that *f* has a *unit-speed parametrization*, which means that ||*f*^*′*^(*t*)||_2_ = 1 for all *t*. Any parametric curve such that *f*^*′*^(*t*) ≠ 0 for all *t* can be reparametrized to satisfy this requirement (Section S1.2). In particular, this means the arc-length between two points on the curve is equal to the difference in their coordinates:

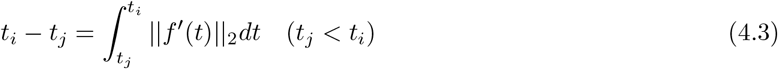

### First morphologically relevant coordinate

Our approach to estimate *t*_*j*_ in the case of a non-intersecting curve (i.e., *t*_1_ ≠ *t*_2_ ⇒ *f* (*t*_1_) ≠ *f* (*t*_2_)) leverages the relationship in (4.3). In fact, our estimated 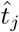 is closely related to the first component produced by the ISOMAP algorithm for manifold learning (Tenenbaum et al., 2000). The key idea in both our approach and ISOMAP is that the distance (geodesic) between two points on a manifold can be well approximated by their shortest path on a *k*-nearest neighbor graph, provided that *k* is sufficiently small to capture the local manifold structure. The set of points traced by *f* (*t*) is a one-dimensional manifold, and the geodesic between two points *t*_*i*_, *t*_*j*_ is exactly the arclength in (4.3). Thus, by defining *G*_*k*_ to be the (connected) *k*-NN graph constructed from the spatial coordinates *x*_1_, …, *x*_*n*_ and 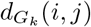 to be the shortest path between *x*_*i*_ and *x*_*j*_ in *G*_*k*_, we have the approximation

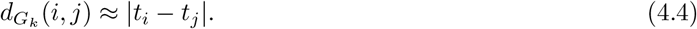

Bernstein et al. (2000) rigorously establishes this approximation under suitable regularity conditions (e.g., uniform sampling along the curve). We demonstrate through simulation (on the swiss roll dataset), that the approximation is very accurate for *n* ≥ 500 (**Fig S20**).

Our final estimate 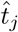 is chosen to satisfy the approximation in (4.4). To construct this, we follow the steps of classical multidimensional scaling (cMDS) (Torgerson, 1952). Squaring both sides of (4.4) yields

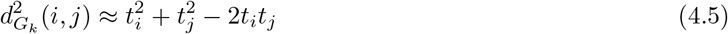

Defining 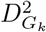 as the *n* × *n* matrix of distances, the operation of double centering (Lemma S1.1) yields 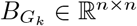 such that

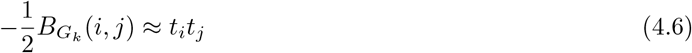

Given this, we set

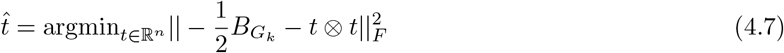

where ⊗ is the outer product (defined in Section S1.1). The optimization problem in (4.7) has (under mild conditions) a closed form solution given by the leading eigenvector of 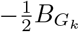 (scaled by the square root of the leading eigenvalue), see Lemma S1.2. In practice, we standardize the resulting 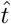 so that it takes values between 0 and 1.

Once 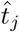 is obtained, the curve *f* can be estimated by smoothing each component function separately. We plug in 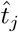 to (2.1) to obtain

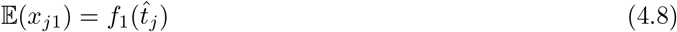

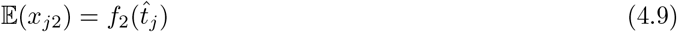

We obtain 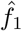 and 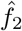 by using penalized regression splines as implemented in *mgcv* (Wood, 2017).

### Extending the method to closed curves

For closed curves, we have *f* (*a*) = *f* (*b*), violating the non-intersecting condition required above. In this case, the approximation in (4.3) no longer holds because there could be a shorter path passing over the endpoint. However, we can still obtain an explicit form for 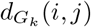, the *k*-NN graph geodesic between cells *i* and *j*. In Section S1.4, we derive in detail the following approximation based on the law of cosines:

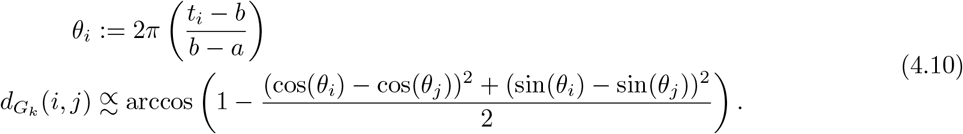

In the above, 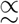 indicates that 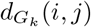 is approximately proportional to the quantity on the right-hand side. We then make a second order Taylor approximation to 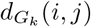:

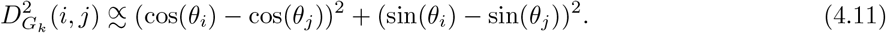

As in the case of an open curve, we apply the double centering operation, yielding

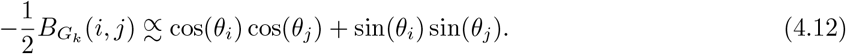

The approximation in Equation (4.12) suggests that 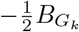 is approximately rank 2 and that *θ*_*i*_ may be recovered by

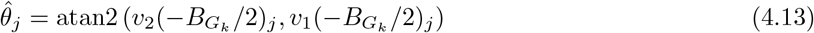

where *v*_*k*_(·) denotes the *k*-th leading eigenvector of a matrix and atan2 is the 2-argument arctangent function (see Section S1.4 for a detailed explanation of this claim). Thus 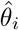 can be converted back to 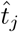 via equation (4.13), although our downstream analysis of SVG detection will be invariant to this scaling.

### Second morphologically relevant coordinate

The second morphologically relevant coordinate 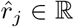 describes how far from the estimated curve a cell’s coordinates are. The magnitude of the coordinate is defined as

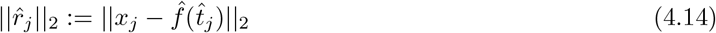

The sign of the second coordinate is determined by

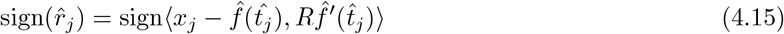

where *R* : ℝ^2^ → ℝ^2^ is a counter-clockwise rotation by 90 degrees: *R*(*v*_1_, *v*_2_) = (−*v*_2_, *v*_1_).

The intuition behind this equation is that cells/spots with a positive sign would be on the left-hand side if one was driving along the curve. The left-hand side is identified by a counter-clockwise rotation *R* of the velocity vector *f*^*′*^(*t*). Again, in practice we standardize 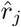 to be in the interval [0, 1] although this could be modified depending on the specific scientific question.

### Selecting k

We provide an automatic approach to selecting the number of nearest neighbors *k*, although we note that we have found the fitted curve and downstream SVGs are typically robust to moderate differences in *k* (**Fig S14**). If *k* is chosen too large, the shortest paths in the graph can skip over the local manifold structure. On the other hand, if *k* is too small the resulting coordinates 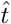 can be very noisy. We develop a heuristic *curve score* that aims to select an ideal *k* that avoids both of these issues.

The score is based on the commonly used Akaike information criterion (AIC) (Akaike, 1974) for choosing the effective degrees of freedom in smoothing splines. Specifically, we let 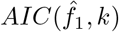 denote the AIC corresponding to the 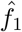 obtained from using *k*, see S1.5 for the full definition. However, the AIC alone does not account for the noise in 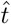. For this reason, we also add a penalty of 2*n/k*, which is related to the effective number of disjoint neighborhoods. The final score becomes:

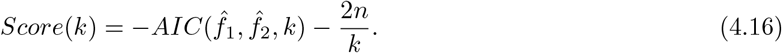

The best choice of *k* can be identified by performing a grid search (by default, we search 3 ≤ *k* ≤ 20) and selecting the value with the largest *Score*(*k*).

### Manual specification of *f* (*t*)

In some datasets, such as the Swiss rolls in Figure 6a, the high density of the samples leads to a *k*-NN graph that is unable to adequately approximate the geodesic distances. Moreover, users may sometimes be interested in identifying variation along a path that does not necessarily correspond to morphology. In these cases, MorphoGAM permits manual specification of the curve *f* (*t*). Using our interactive software, the user selects points *t*^(*k*)^ = (*x*^(*k*)^, *y*^(*k*)^), 1 ≤ *k* ≤ *K* which are then smoothed (using a GAM) to obtain estimates of *f* . Given this *f, t* is estimated using the projection index of Hastie and Stuetzle (1989):

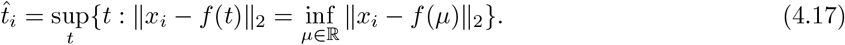

Estimation of *r* then proceeds as before.

### Checking the one-dimensional assumption

In situations where MorphoGAM is applied to many different tissue samples, it can be useful to have a diagnostic to check the validity of the one-dimensional manifold assumption. When the assumption is valid, the arclength of the fitted curve will be expected to be much larger than the distance of a typical cell to its projection onto the curve. This motivates the definition of our *ratio diagnostic RD*, which is the arclength (as defined in (4.2)) divided by the 0.95-quantile of 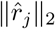 (before standardizing, as defined in (4.14)). We computed the *RD* for three of the examples with clear one-dimensional structure and observed that *RD* ranged between 8 and 35. On the other hand, when we applied MorphoGAM to points drawn from the unit square, we observed *RD* ≈ 1.8 (Table S1), indicating that the *t* and *r* dimensions have similar scales. Values much smaller than 8 could therefore warrant manual inspection to decide if MorphoGAM is a suitable tool.

### Disconnected graphs

The estimation procedure described above requires *G*_*k*_ to be a connected graph. However, if *k* is chosen large enough to ensure the graph is fully connected, then 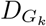 may not capture more subtle morphological features. For this reason, we permit the procedure to be applied separately to disconnected components of *G*_*k*_ and then *stitched* together to create the final curve. Given *G*_*k*_ has *C* connected components, let 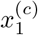 and 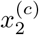 denote endpoints of the curve describing the *c*-th component. We then identify the connections between 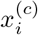 and 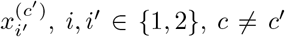 of minimum Euclidean distance that produce a single (connected) curve. Note that this may require reversing the direction of a curve fit to one particular component. We identify the optimal connections through a brute force search of the *C*! · 2^*C*^ possibilities. Because this is computationally infeasible for large *C*, we require that *k* is at least large enough to ensure that *C* ≤ *C*_max_, where *C*_max_ = 5 by default. Note also that outlier points can lead to a disconnected graph, and these can optionally be removed by the MorphoGAM procedure.

### Generalized additive model to identify spatially variable genes

Let *Y*_*gj*_ denote the count for gene *g* (1 ≤ *g* ≤ *G*) in cell/spot *j* (1 ≤ *j* ≤ *n*). As noted before, we consider the model

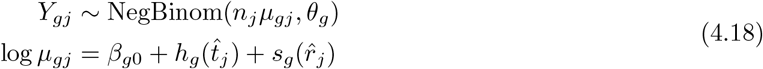

where *h*_*g*_ and *s*_*g*_ are unknown smooth functions, *β*_*g*0_ is an unknown intercept, and 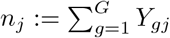 is a known offset. Note that we are using the following standard parameterization of the negative binomial distribution:

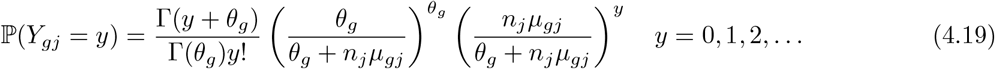

For identifiability, we also assume that 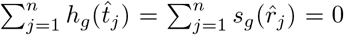. We use *mgcv* (Wood, 2017) to estimate the functions in model (2.6); this method writes *h*_*g*_ and *s*_*g*_ as a linear combination of known basis functions 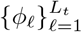 and 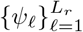

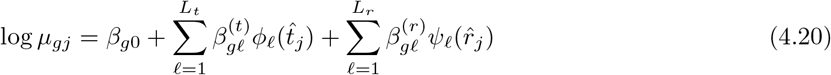

where *L*_*r*_ and *L*_*t*_ are sufficiently large (*L*_*r*_ = *L*_*t*_ = 10 by default). Although there is flexibility in the choice of *ϕ* and *ψ*, we use cubic regression splines (cyclic for *ϕ* when *f* is a closed curve). Estimation proceeds by maximizing the log-likelihood of the model parameters 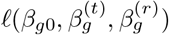 (here 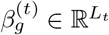 and 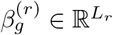 are vectors of coefficients) subject to a smoothness penalty:

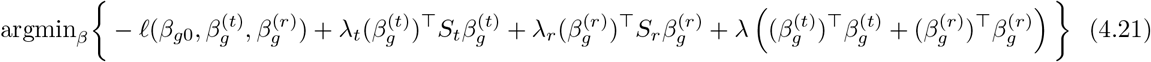

where *S*_*t*_ and *S*_*r*_ are (known) matrices depending on the second derivative of the chosen basis functions (Wood, 2001). *mgcv* performs a procedure (generalized cross validation, (Golub et al., 1979)) to select the best choice of *λ*_*t*_ and *λ*_*r*_ and we set *λ* = 1 by default. Upon estimating the coefficients, *mgcv* returns a posterior covariance matrix for uncertainty quantification. Because the fitted functions for each genes can have different amounts of uncertainty, we shrink the functions (pointwise) by amount based on its standard error prior to computing the peak and range functionals. In particular, if *ĥ*(*t*) is positive we set 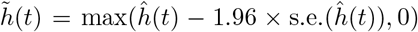 and 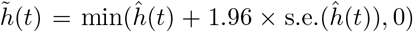 otherwise. These values are chosen because this will set 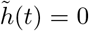 if the 95% confidence interval around *ĥ*(*t*) contains 0. For ranking genes, we have found this hard-thresholding to be useful as only the most relevant genes will typically obtain non-zero values. On the other hand, for visual inspection of 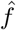 (and for the SVD analysis to find groups of genes), we instead opt to use a smooth shrinkage that prevents *f* from being exactly 0. In particular, we set 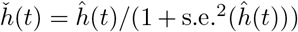. This corresponds to the posterior mean of *h*(*t*) under a standard normal prior.

### Identifying genes with correlated expression patterns

To identify the dominant patterns of spatial variability, we aim to decompose each function *ĥ*_*g*_ into a sum of *M* functions *ψ*_*i*_:

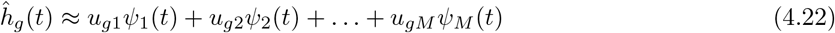

This decomposition can be obtained using a rank *M* singular value decomposition (SVD) of the matrix *Ĥ* ∈ ℝ^*G×n*^ with row *g* corresponding to *ĥ*_*g*_ (*t*). Because the function estimates from the GAM are already smooth, the right singular vectors of *Ĥ* can directly be used as an estimate of *ψ*_*m*_. The exact same approach can be applied to the *r*-direction functions *ŝ*_*g*_ as well. We note that this procedure has similarities to functional principal component analysis (FPCA) (Wang et al., 2016).

### Details of simulation study

Using the spatial coordinates from the CA3 cells (*n* = 1456) we simulated *G* = 1000 genes from a model with *Y*_*ig*_ ∼ NegBinom(*µ*_*gj*_, 10). For gene 1, we set 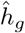 and considered three patterns of spatial variability for the function *h*:

- *Localized:* We constructed a natural spline basis *B*_1_(*t*), …, *B*_10_(*t*) on the interval [0, 1] using ns. We then selected a basis function *B*_*k*_(*t*) (uniformly at random) and set *h*(*t*) = *B*_*k*_(*t*). This creates a pattern with a localized peak, as shown in Figure 4d.
- *Gradient: h*(*t*) = *t*.
- *Non-parametric:* 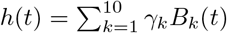, where *γ*_*k*_ ∼ *N*(0, 1). By scaling, we ensured that sup_*t*_ |*h*(*t*)| =1.

For the *r*-associated genes 2 through *G*, we set 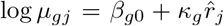. We drew 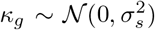. The three scenarios low *r*, medium *r*, high *r* correspond to *σ*_*s*_ = 0.5, *σ*_*s*_ = 1, *σ*_*s*_ = 2, respectively. We sampled *β*_*g*0_ ∼ N(−1, 1). For each pattern and scenario, we performed 10 independent replicates.

We obtained a ranked list of genes from each of the methods using the following approach:

- *SPARK-X* (Zhu et al., 2021), *HEARTSVG* (Yuan et al., 2024): The rank was determined using the adjusted *p*-value for each gene.
- *nnSVG* (Weber et al., 2023): A rank column is directly outputted.
- *GASTON* (Chitra et al., 2025): Setting the number of domains equal to 1, GASTON fits a piecewise linear function to each gene and produces a *p*-value that was used to rank the genes.
- *Spateo* (Qiu et al., 2024): We used their digitization function (which required manual input) to approximate the *t* and *r* coordinate. We then used their glm degs function with a full model including natural spline terms for both *t* and *r* and a reduced model containing just the term for *r*. The *p*-value was obtained using the likelihood ratio test.
- *MorphoGAM* : Each gene is assigned a peak statistic 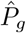 in the *t* direction, which was used to construct the ranking.

For the granule cells (**Fig S12**), the same procedure as above was used to generate simulated counts starting from the (*x, y*) coordinates of the granule cells and using the *t* and *r* coordinates from the reference curve in Figure S3.

## Data and code availability

The following datasets were used:

- The granule cells in **Figure 3** were obtained from the data provided by Cable et al. (2022b). Cells such that the 5-th nearest neighbor was 2 times greater than the median 5-th nearest neighbor were excluded. This procedure removed outlier cells.
- The CA3 cells were obtained from *STexampleData* (Righelli et al., 2022). Cells such that the 20-th nearest neighbor was 3 times greater than the median 20-th nearest neighbor were excluded. These values were used so that the retained set of cells visually matched Figure 5 of Cable et al. (2022b).
- MERFISH measurements of the adult healthy colon are described in (Xu et al., 2026). Briefly, these measurements were performed using standard MERFISH protocols, targeting a custom set of 1,920 genes. The data can be accessed at https://doi.org/10.5061/dryad.p5hqbzm0z.
- The MERFISH swiss rolls of the mouse colitis model are described by (Cadinu et al., 2024) and can be accessed at https://datadryad.org/dataset/doi:10.5061/dryad.rjdfn2zh3.
- The processed Xenium mouse brain was downloaded from the 10X genomics website https://www.10xgenomics.com/datasets/fresh-frozen-mouse-brain-replicates-1-standard. The cell type labels were extracted from the Figure 5a source data of (Fu et al., 2024). Outlier cells were removed prior to curve estimation.
- The Visium dorsolateral prefrontal cortex were originally generated by Maynard et al. (2021) and obtained from *STExampleData* Righelli et al. (2022).
- The Visium zebrafish melanoma data and H&E images (Hunter et al., 2021) are available on GEO with accession number GSE159709.

MorphoGAM is available as an R package at https://github.com/phillipnicol/MorphoGAM. The repository https://github.com/phillipnicol/morphoGAM-paper includes scripts to reproduce all results in the paper.

## Acknowledgements

PBN was supported by the National Institutes of Health grant T32CA009337. JRM was supported by the National Institutes of Health grant R01GM143277.

## Supplementary Material

### S1 Mathematical details of curve and coordinate estimation

#### S1.1 Notation

We use ⊗ to denote outer-product: if *u, v* ∈ ℝ^*n*^ then *u* ⊗ *v* := *uv*^⊤^ ∈ ℝ^*n×n*^. If *f* : [*a, b*] → ℝ^*k*^ denotes a parametric curve, then *f* can be written in terms of *k component functions f* (*t*) = (*f*_1_(*t*), …, *f*_*k*_(*t*)) and 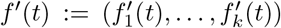. We say *f* is *smooth* if 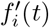 exists and is continuous for all *t*. For a matrix 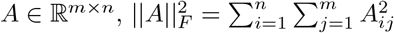 denotes Frobenius norm.

#### S1.2 Assumptions

We assume the following conditions, which are necessary (but not sufficient) for the identifiability of model (2.1):

1. 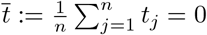 and *t*_1_ *<* 0.

2. ||*f*^*′*^(*t*)||_2_ = 1 for all *t*.

For condition 1, note that for any constant 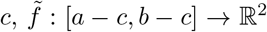 defined by 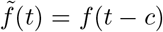 satisfies 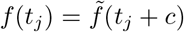 for every *j*. Similarly, 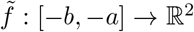 defined by 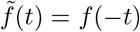 satisfies 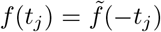. For condition 2, define 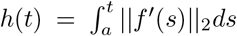 and note that *h*^*′*^(*t*) = ||*f*^*′*^(*t*)||_2_ . Then we can define the reparameterized curve 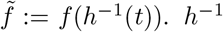 exists and is differentiable by the inverse function theorem. In particular,

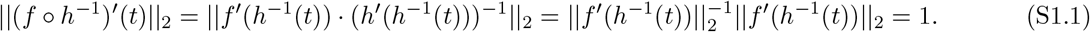

A full introduction to parametric curves is given by Lastra (2021). We also note that these conditions are not sufficient to ensure the identifiability of model (2.1) as a fully identifiable model would likely need to specify a distribution or a procedure from which the *t*_*j*_ are obtained.

#### S1.3 Open curve

We now describe how to estimate the first coordinate in the case of an open, non-intersecting, curve (i.e., *t*_1_≠ *t*_2_ ⇒ *f* (*t*_1_) ≠ *f* (*t*_2_)). If we assume that the approximation in equation (4.4) is equality, i.e., 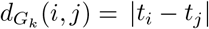, then 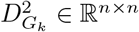 can be written in matrix form as

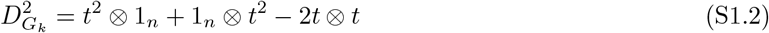

where *t*^2^ is applied entry-wise to *t* := (*t*_1_, …, *t*_*n*_) and 1_*n*_ ∈ ℝ^*n*^ is a vector of 1’s. Now define the centering matrix *H* ∈ ℝ^*n×n*^ as

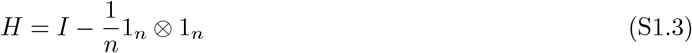

Applying *H* on the right has the property of subtracting the row means while applying *H* on the left subtracts the column means.

**Lemma S1.1**. *The “double centered” matrix* 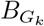 *satisfies*

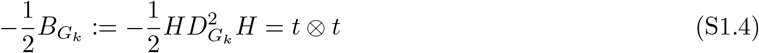

*Proof*. The proof is derived from Ghojogh et al. (2023). Because *H*(1_*n*_ ⊗ *t*^2^) = 0 and (*t*^2^ ⊗ 1_*n*_)*H* = 0, we have

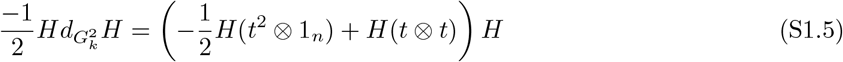

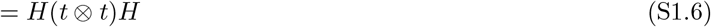

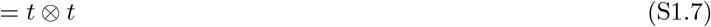

where the last line follows because 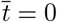 by assumption.

The above result shows that 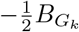 is a rank 1 matrix with a positive eigenvalue ⟨*t, t*⟩ *>* 0. In practice, however, 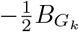 will be expected to have higher rank due to noise. For this reason, we estimate *t* using the top eigenvector associated with the largest eigenvalue. The following lemma shows that, under some conditions, the top eigenvector of a symmetric matrix leads to the best rank-one approximation with the smallest reconstruction error. The result is a simple application of low-rank approximations (Eckart and Young, 1936).

##### Lemma S1.2

*Let A* ∈ ℝ^*n×n*^ *be a symmetric matrix, and suppose that λ*_max_(*A*) *>* 0 *and λ*_max_(*A*) *>* |*λ*_min_(*A*)|, *where λ*_max_ *and λ*_min_ *denotes the largest and smallest eigenvalues, respectively. Then*

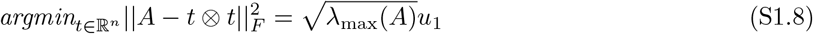

*where u*_1_ *is the unit eigenvector corresponding to λ*_max_(*A*).

*Proof*. As *A* is symmetric, we may write

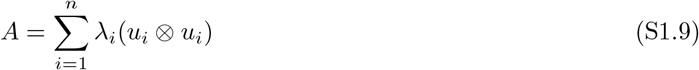

with *u*_1_, …, *u*_*n*_ ∈ ℝ^*n*^ orthonormal. Then

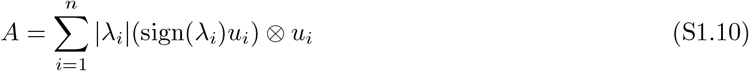

is a singular value decomposition (SVD) of *A*. By Eckart and Young (1936), we have

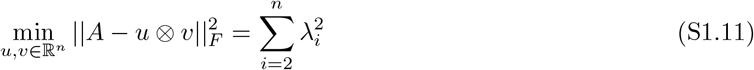

Moreover,

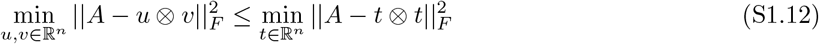

so 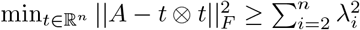 as well. This minimum can be achieved by setting 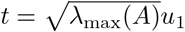, as *λ*_max_(*A*) = *λ*_1_.

In practice, it seems to be the case that the conditions *λ*_max_(*A*) *>* 0 and *λ*_max_(*A*) *>* |*λ*_min_(*A*)| hold.

#### S1.4 Closed curve

In the case of a closed curve *f* (*a*) = *f* (*b*), it is useful to first transform the domain to the unit circle *S*^1^ by writing 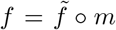 where *m* : [*a, b*] → *S*^1^ maps *t* to (cos(2*π*(*t* − *b*)*/*(*b* − *a*)), sin(2*π*(*t* − *b*)*/*(*b* − *a*)). For convenience, we will write *θ*_*i*_ = 2*π*(*t*_*i*_ − *b*)*/*(*b* − *a*). Because *f* is unit speed,

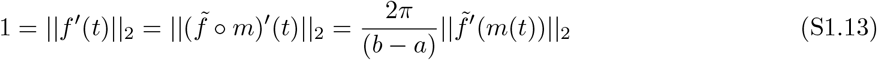

Showing that the curve 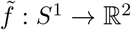 is constant speed and that 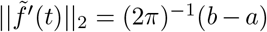. The shortest path between cells *i* and *j* in the *k*-NN graph *G*_*k*_, 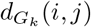, will be approximately equal to the smaller of the two arclengths connecting 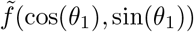 and 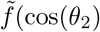, sin(*θ*_2_)). Because 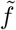 is constant speed, this arclength will be equal to (2*π*)^−1^(*b* − *a*) times the (shorter) arclength between (cos(*θ*_1_), sin(*θ*_1_)) and (cos(*θ*_2_), sin(*θ*_2_)) on the unit circle. This may be computed explicitly by defining *δ* to be the Euclidean distance between (cos(*θ*_1_), sin(*θ*_1_)) and (cos(*θ*_2_), sin(*θ*_2_)) and using the law of cosines:

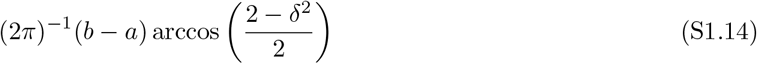

Expanding *δ*^2^ and noting that (2*π*)^−1^(*b* − *a*) is a constant yields the approximation stated in equation (4.12).

Because (S1.14) appears to have no simple form, we make a second order Taylor approximation (as a function of *δ*). Using the power series representation of arccos (DLMF, (4.24.2)) gives

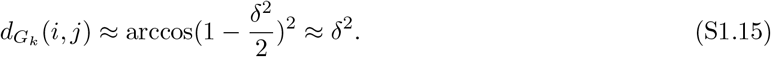

By an argument similar to Lemma S1.1, applying the double centering operation to 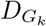 yields

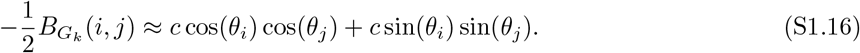

for some constant *c*. This implies that 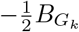 will be approximately rank 2. Furthermore, if *n* is large and *θ*_*i*_ densely populated within [0, 2*π*] then we have

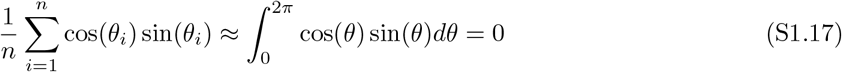

which shows that (S1.16) is also an (approximate) eigendecomposition of 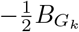. In particular, if the two eigenvectors recover cos(*θ*_*i*_) and sin(*θ*_*i*_), respectively, then taking the arctangent function of the ratio should be a reasonable approximation to *θ*_*i*_. Because both cos(*θ*) and sin(*θ*) are approximate eigenvectors with eigenvalue *c*, the top two eigenvectors could be invariant to rotation. However, in any case, the top two eigenvectors would still represent the location of each cell on a circle, and the two-argument arctangent function would still recover the angle along that circle.

#### S1.5 Details of curve score

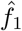 and 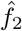 are obtained using *mgcv*, which assumes the following model

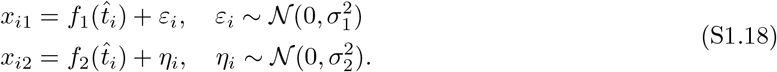

Using the above model, the log likelihoods 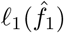 and 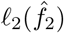 can be defined for any estimate 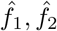 of *f*_1_, *f*_2_. The regression spline procedure implemented by *mgcv* produces hat matrices *H*_1_, *H*_2_ ∈ ℝ^*n×n*^ such that 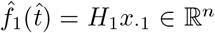 and 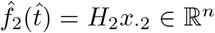. The *effective degrees of freedom* for *f* 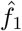 (given 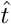) is defined as the trace of *H*_1_, Tr(*H*_1_). Plugging this value into the definition of AIC gives

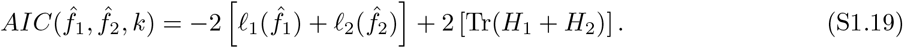

Note that the estimators implicitly depend on *k* through 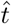.

If the (effective) number of parameters required to estimate 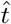 was exactly *n/k*, then the AIC of the entire model used to estimate 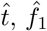, and 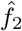 would be *Score*(*k*) (4.16). We provide heuristic justification for this. For a given value of *k*, there are ≈ *n/k* disjoint neighborhoods. Within each neighborhood, one could define a landmark point. Then the shortest paths between cells in the graph are well approximated by the shortest paths between the corresponding landmarks. Thus ≈ *n/k* parameters suffice to produce a good approximation to 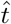.

### S2 Technical details of simulations and results

#### S2.1 Statistical power analysis

MorphoGAM can increase statistical power to detect trajectory-specific SVGs by projecting the two-dimensional ST coordinates to a one-dimensional morphologically relevant coordinate. To demonstrate this, we simulated a gene such that 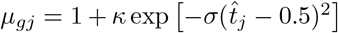, where 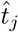 is as above. To compare to the approaches based on hypothesis testing, we labeled a gene as spatially variable if the *p*-value was below the transcriptome-wide significance level of 0.05*/*20000. MorphoGAM had consistently higher power than nnSVG (Weber et al., 2023) and SPARK-X (Zhu et al., 2021) (**Fig S18**). To show that the increase in power did not come at the price of an inflated type I error rate, we randomly permuted all spatial locations to generate a null dataset with no spatially variable genes, and found that our method was conservative at a variety of significance levels (**Fig S19**).

#### S2.2 Testing for markers of inflammation-associated fibroblasts

Recall the model (2.9) for identifying markers of IAFs in a single swiss roll:

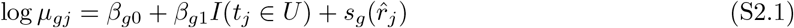

where *I*(*t*_*j*_ ∈ *U*) = 1 if cell *j* is in the ulcerated region and is 0 otherwise. We fit this for the three swiss rolls, obtaining 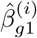, where *i* ∈ {1, 2, 3}. To identify positive markers of IAFs, we define 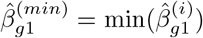. Genes with a large value of 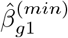 have persistent signal across all three swiss rolls. To obtain a *p*-value, first define *p*^(*i*)^ to be the *p*-value corresponding to the test 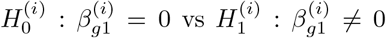. Our null hypothesis is 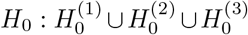. That is, *H*_0_ is true if at least one of the 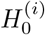 is. A conservative *p*-value is, as (assuming 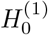 is true):

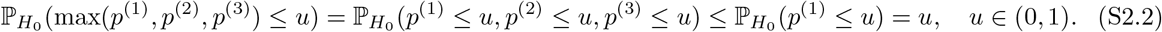

#### S2.3 Details of solid tumor analysis

In the Hunter et al. (2021) analysis, the non-tumor spots are removed by filtering for *r* ≥ 0.5. The *t* coordinate is not meaningful near the tumor core. For this reason, we fit a modification of model (2.6) that only permits edge effects:

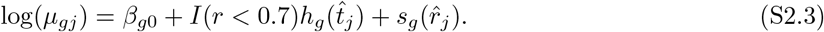

where *h*_*g*_ satisfies *h*_*g*_(0) = *h*_*g*_(1). The above model is specified by the following design formula in R:

`∼ s(t, bs = “cc”, by = edge) + s(r, bs = “cr”)` where edge is a binary vector containing the values of *I*(*r*_*j*_ *<* 0.7).

## S3 Supplementary figures and tables

**Figure S1.**
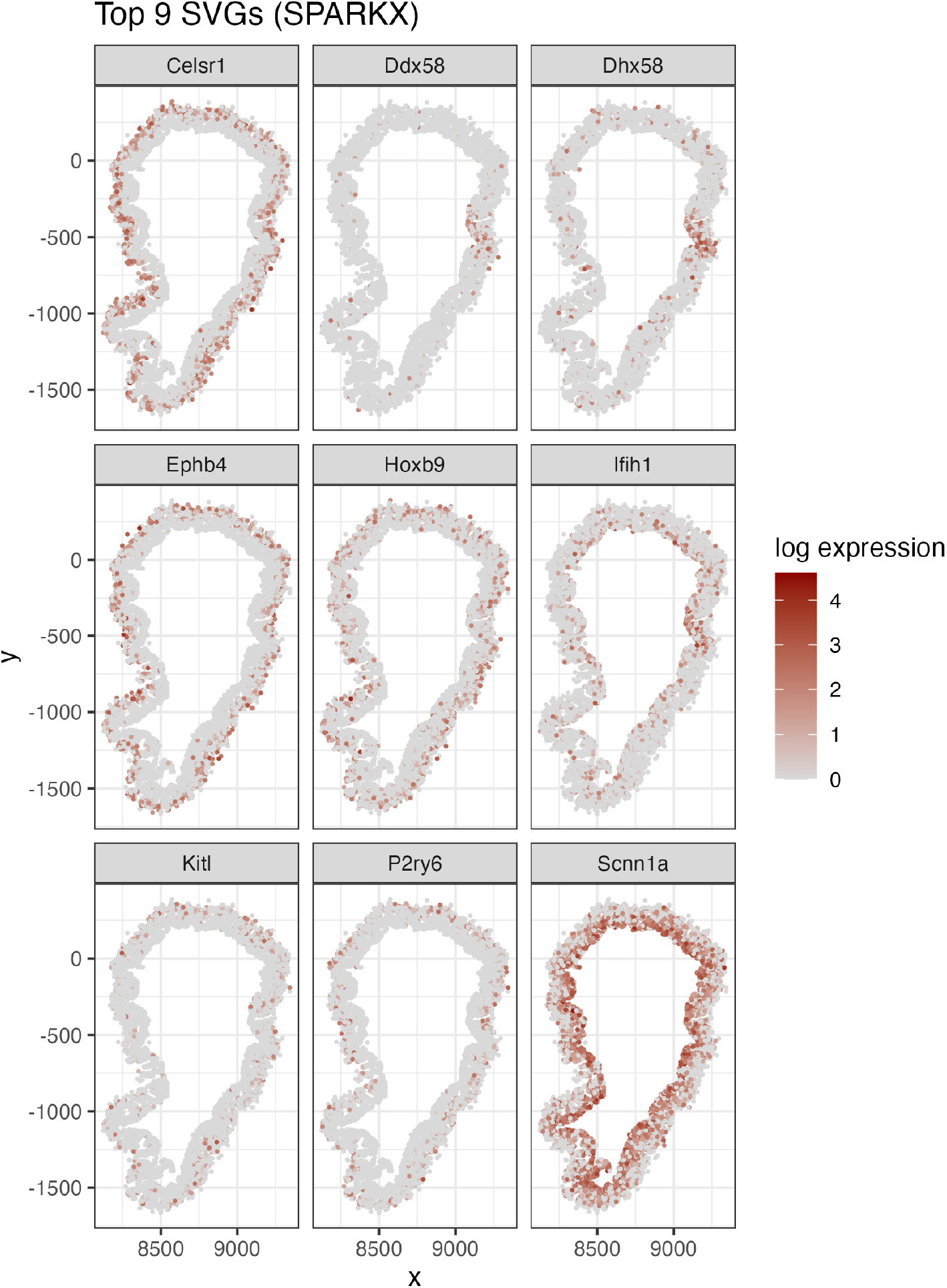
The top 9 SVGs identified by SPARK-X (Zhu et al., 2021) in a MERFISH measurement of a slice of the healthy mouse colon. Specifically, *Ddx58* had a reported (adjusted) *p*-value of 6.38 × 10^−67^ and *Apob* had a reported (adjusted) *p*-value of 8.04 × 10^−9^.

**Figure S2.**
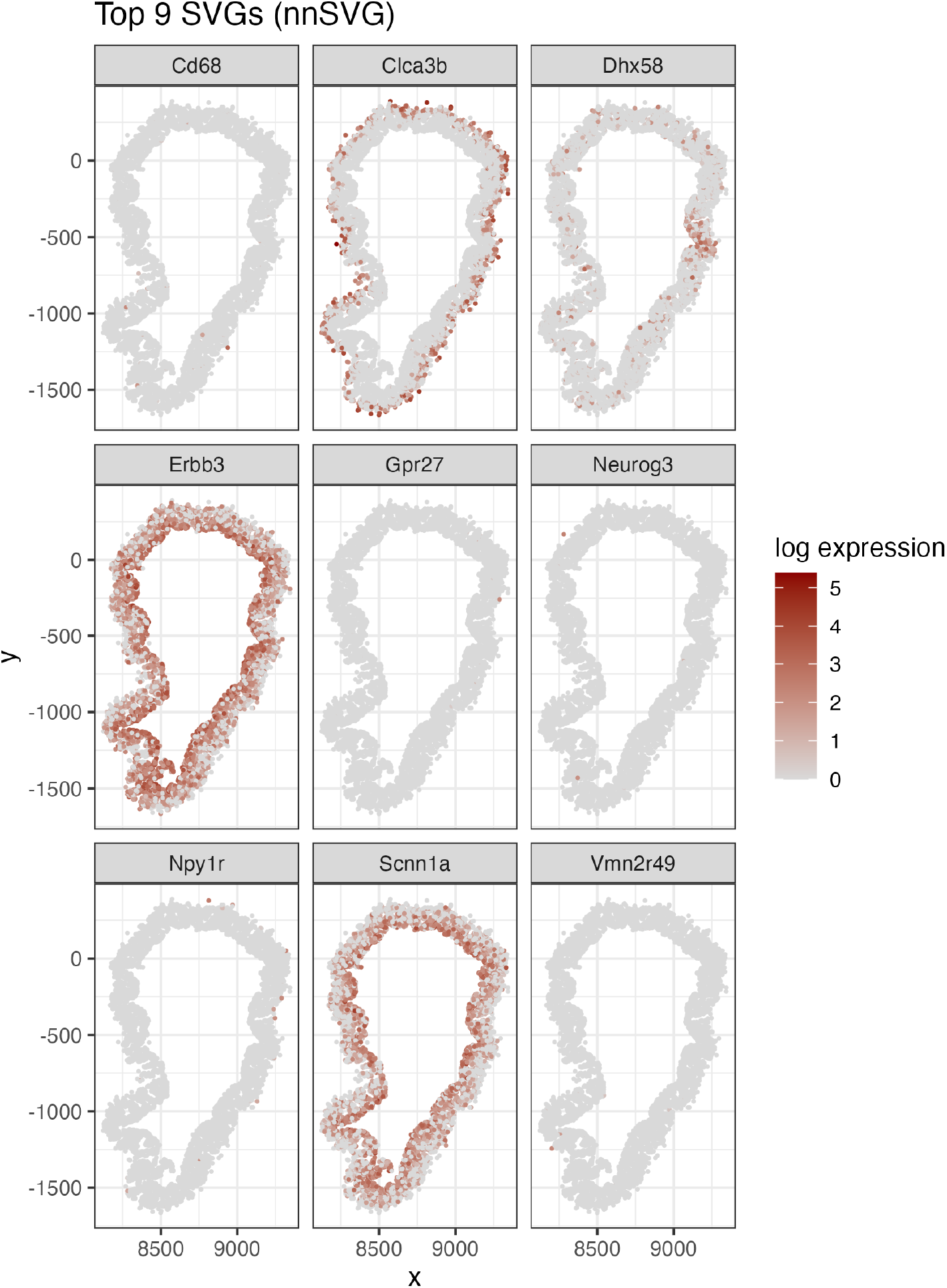
The top 9 SVGs identified by nnSVG (Weber et al., 2023) in a MERFISH measurement of a slice of the healthy mouse colon. Specifically, *Ddx58* and *Apob* both had reported (adjusted) *p*-values of 0.

**Figure S3.**
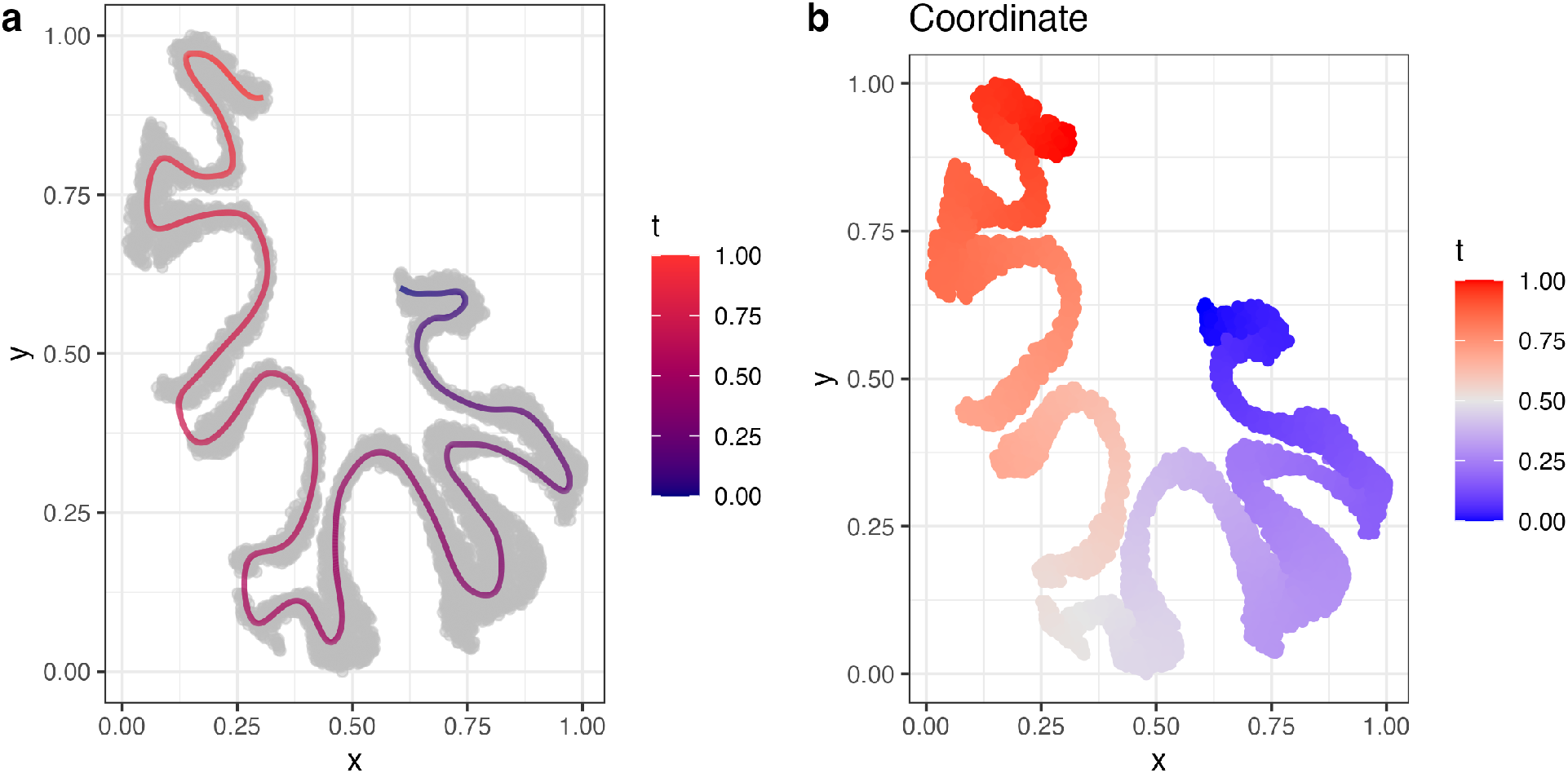
**a**. The hand-drawn curve on the granule cells. **b**. The estimated coordinate from the hand-drawn path on the granule cells.

**Figure S4.**
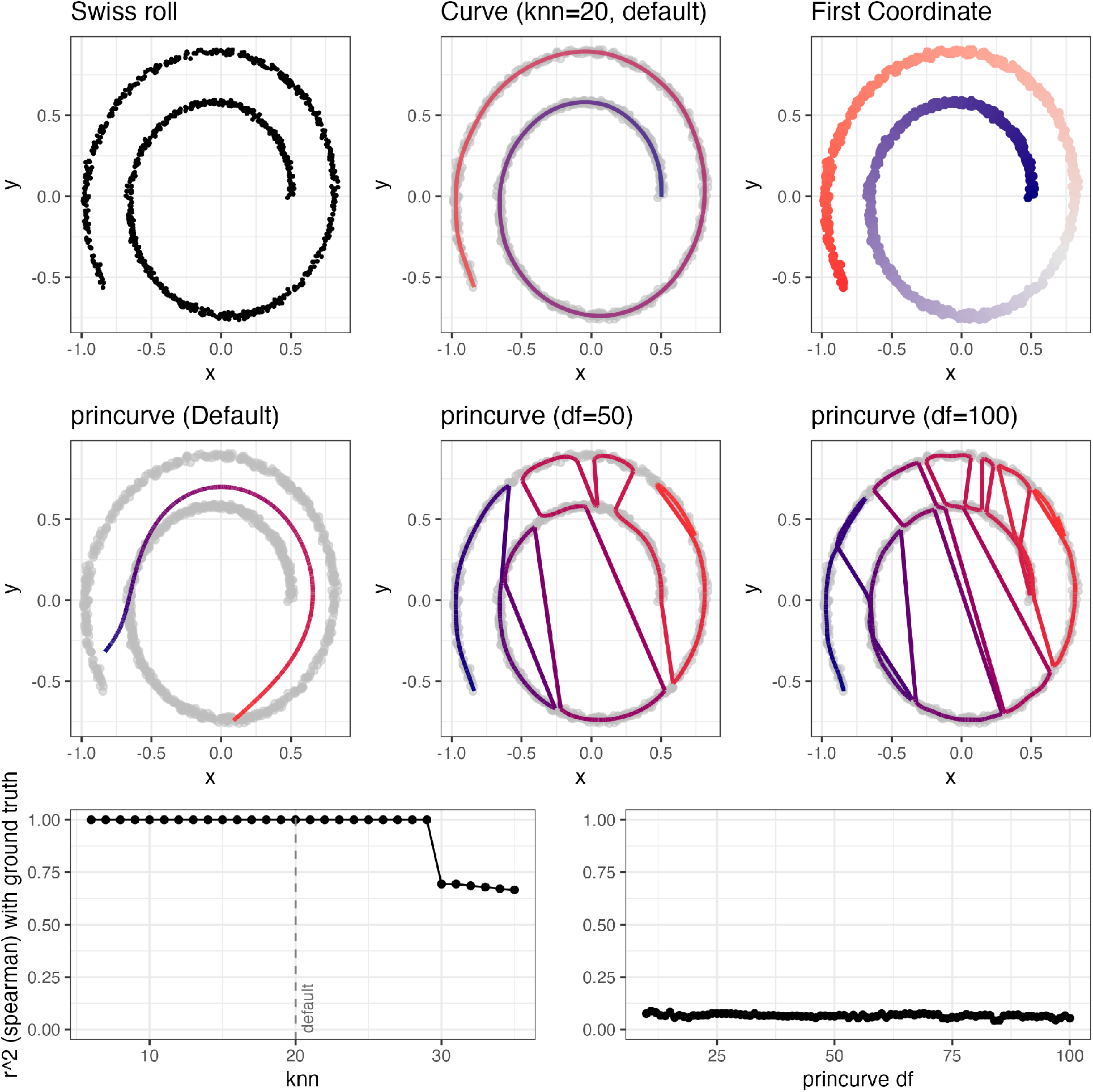
Repeating the analysis in Figure 3 instead using a simulated swiss roll. The inability of the standard principal curves algorithm to accurately reconstruct the swiss roll was discussed in (Kégl et al., 2000).

**Figure S5.**
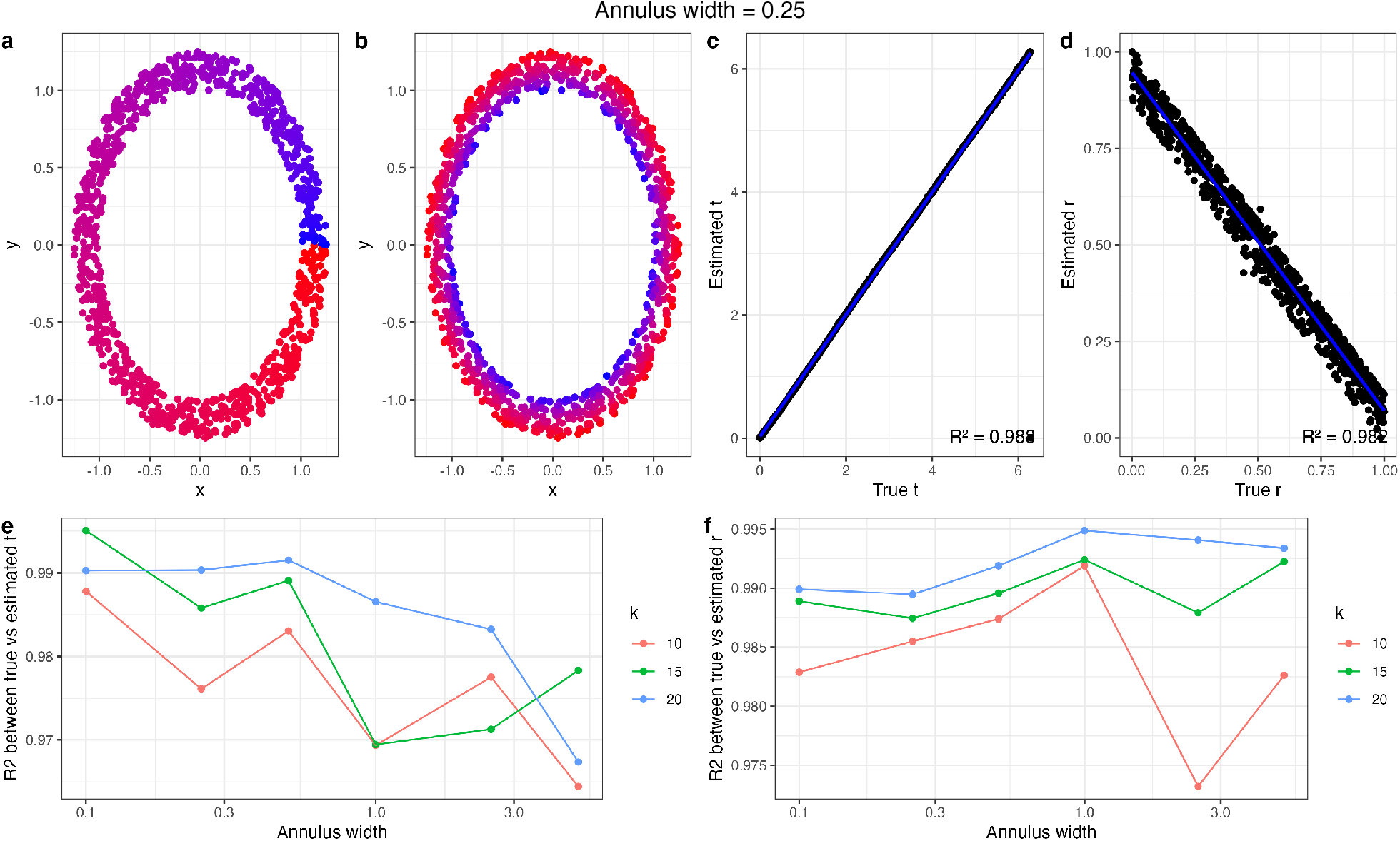
A simulated annulus (width 0.25) dataset to benchmark the estimation of *r*. **a**. The true *t* coordinate. **b**. The true *r* coordinate. **c**. The estimated (by MorphoGAM with *k* = 10) vs true *t* coordinate. **d**. Estimated vs true *r* coefficient. **e**. The width of the annulus and *k* parameter of MorphoGAM is varied, and the *R*^2^ between the true and estimated *t* is reported. Points represent the mean of 10 replicates. **f**. The same experiment, reporting the *R*^2^ between the true and estimated *r* coordinate.

**Figure S6.**
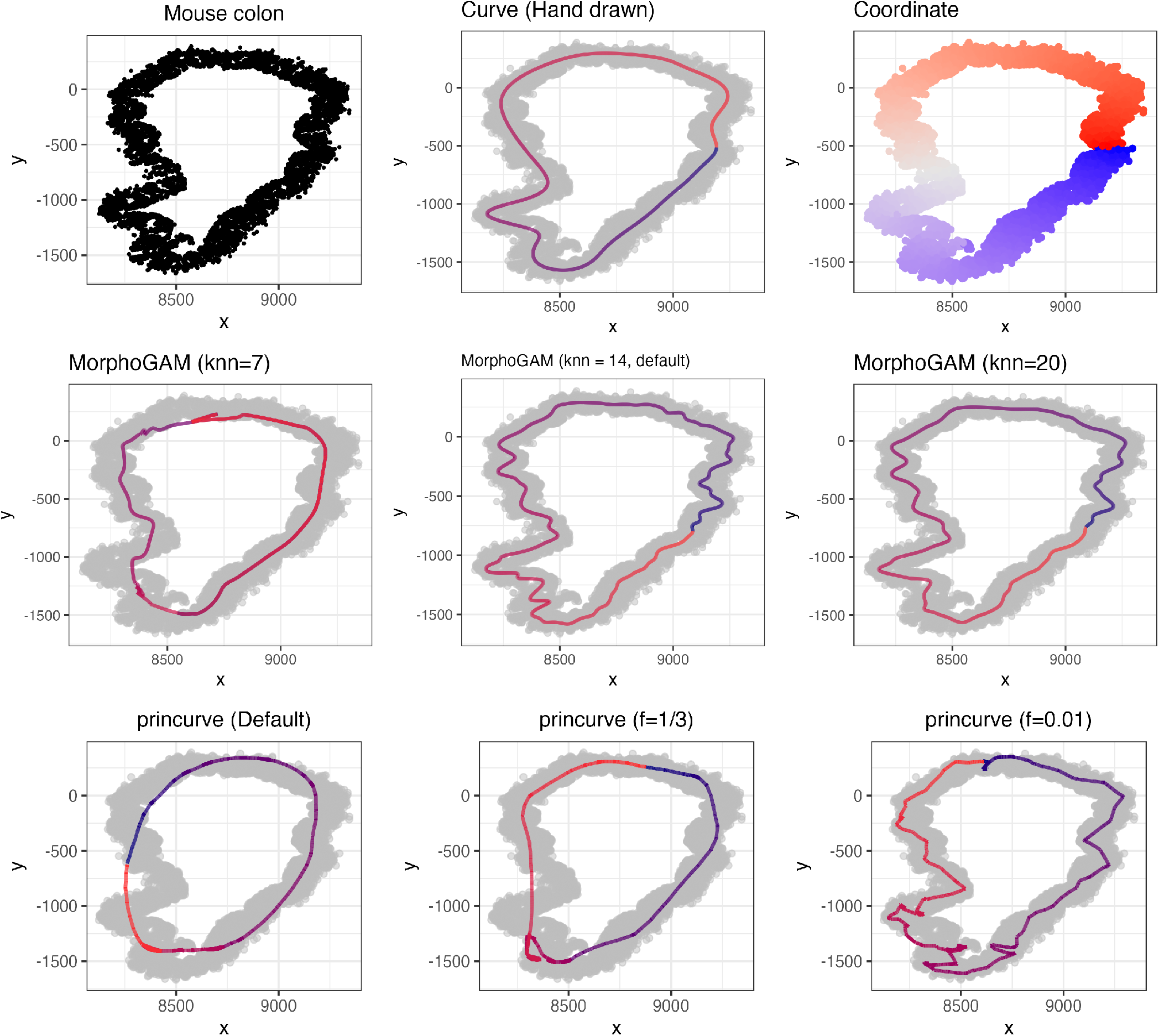
Repeating the analysis in Figure 3 instead using the mouse mucosa data. For MorphoGAM, we demonstrate the default *k* = 14 as well as a value that is too small (*k* = 7). A larger value *k* = 20 yields a slightly smoother curve, yet we show in Figure S14 that the downstream SVGs align very closely with *k* = 14. For principal curves, a periodic smoother was used.

**Figure S7.**
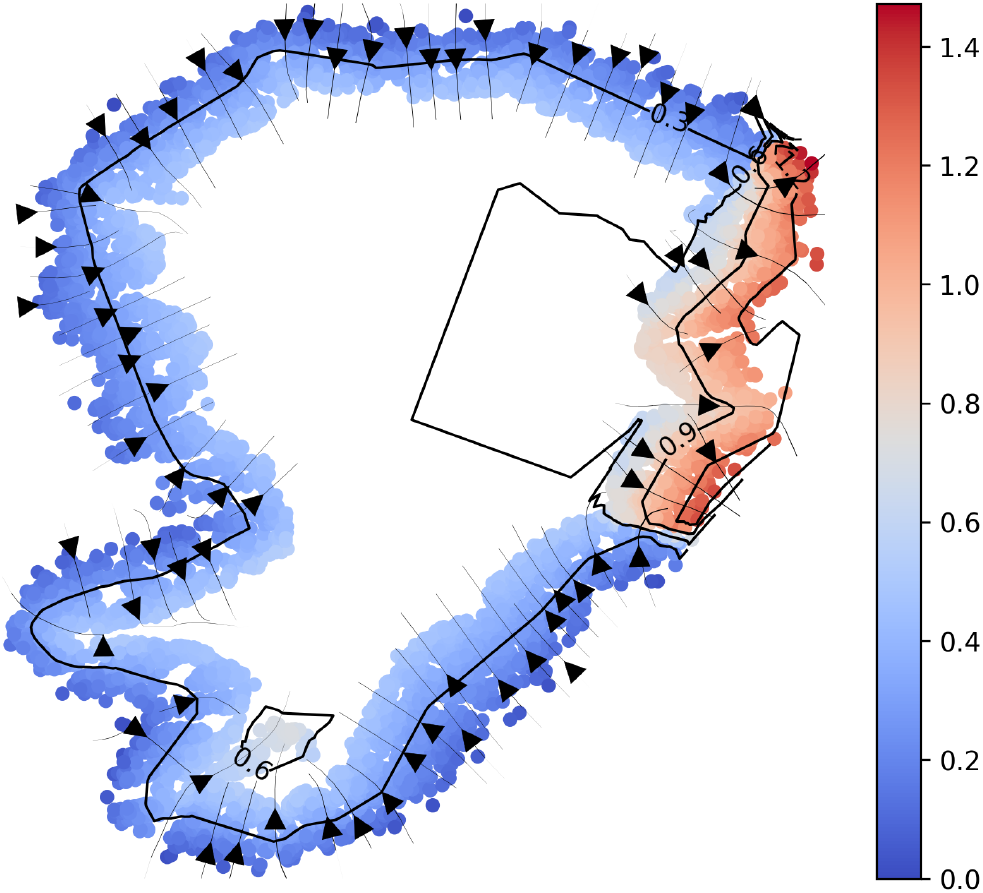
The isodepth obtained by applying GASTON (Chitra et al., 2025) to the MERFISH mouse colon data. Although the isodepth captures the location where *Dhx58* is patchy, the isodepth alone can not be used to distinguish between radial gradient and patchy genes, as is shown in Figure S8.

**Figure S8.**
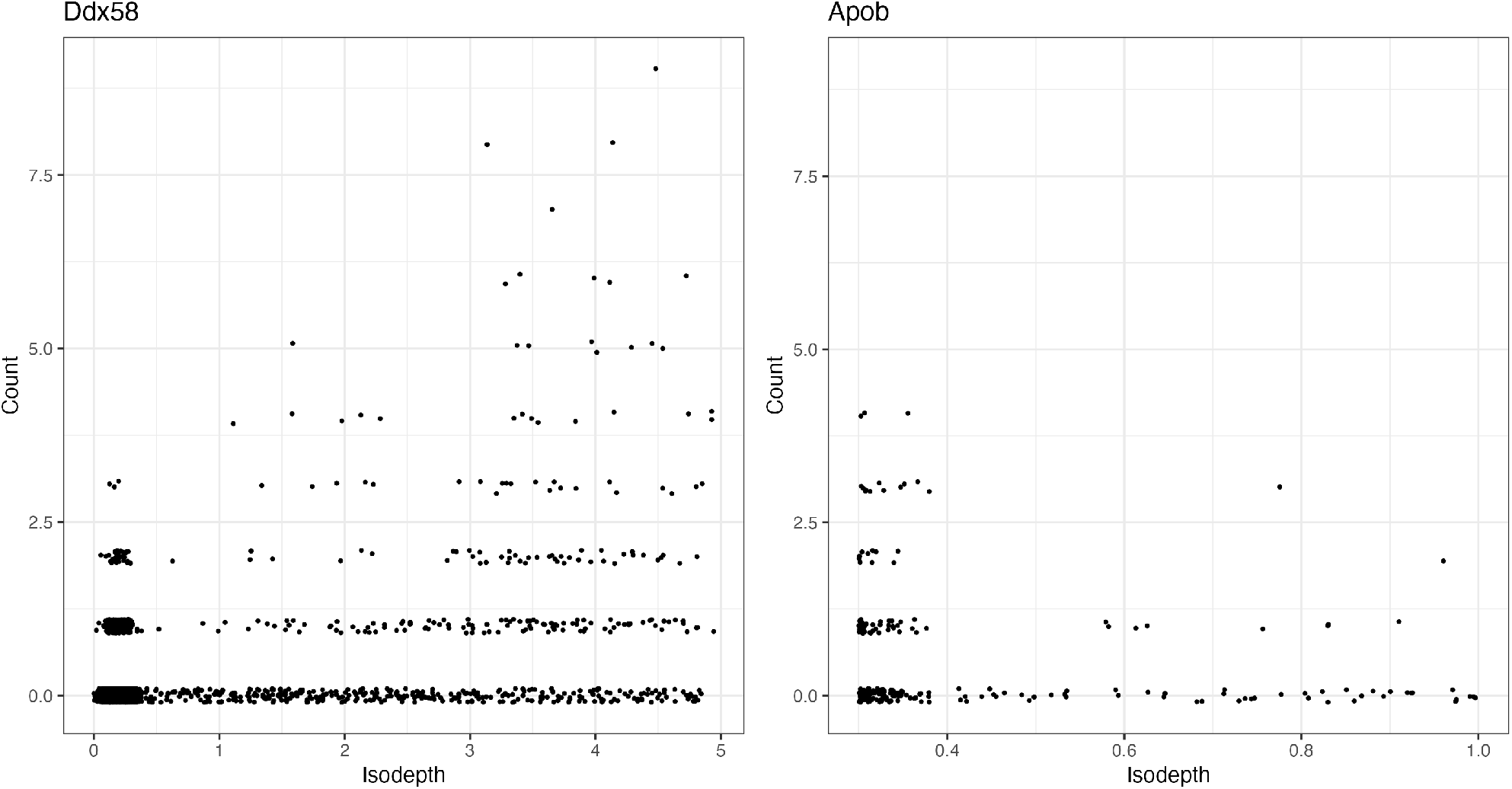
Plotting the two genes of interest (*Ddx58* and *Apob*) from Figure 2 as a function of the isodepth estimated in Figure S7. Both *Ddx58* and *Apob* clearly show variable expression along the isodepth coordinate, showing that this quantity alone can not be used to distinguish between patchy and radial gradient genes.

**Figure S9.**
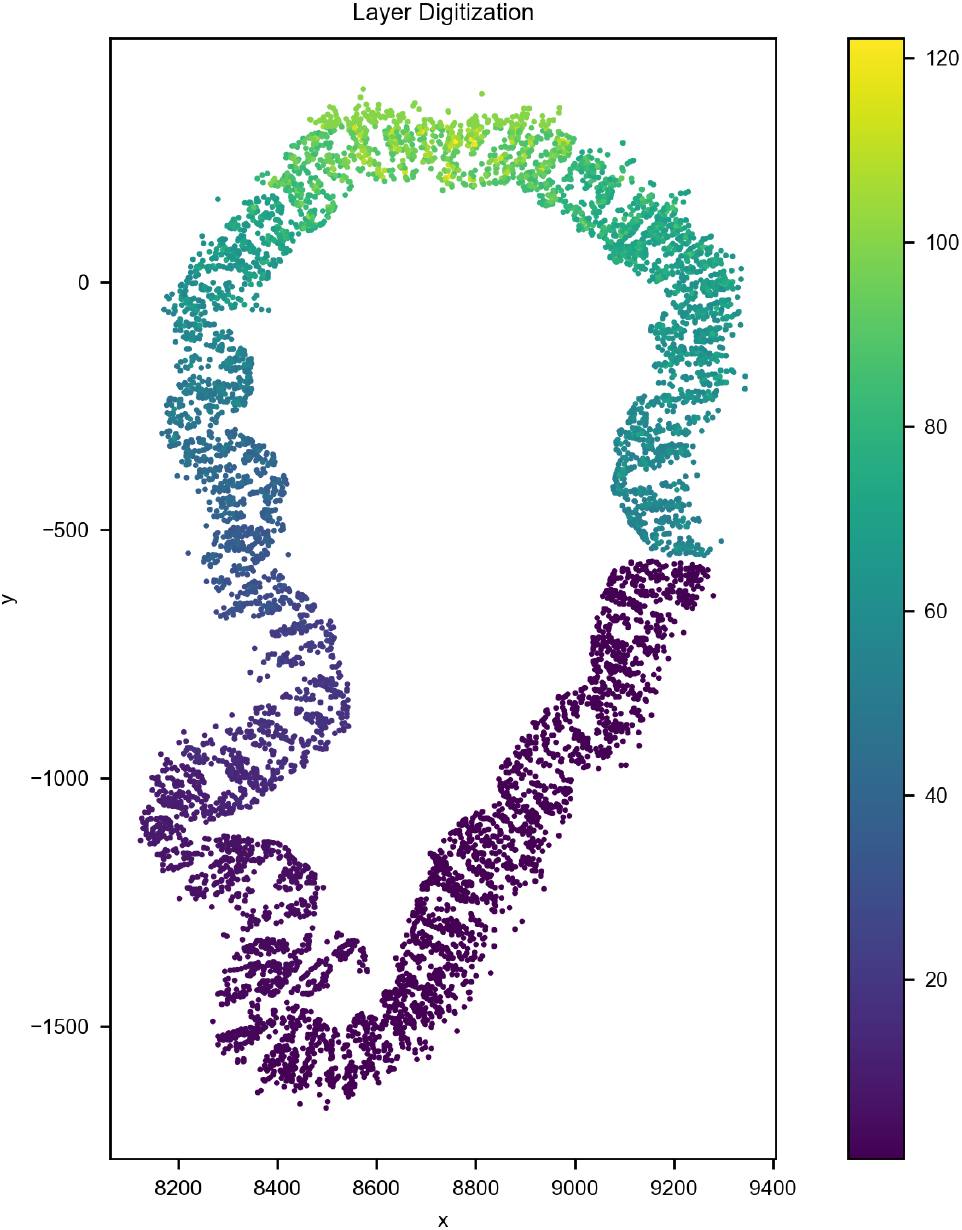
The result of using the Spateo (Qiu et al., 2024) spatial digitization method on the mouse mucosa data. Two points on opposite ends of the tissue are assigned the same coordinate, showing that this approach does not accurately capture the morphology of the colon.

**Figure S10.**
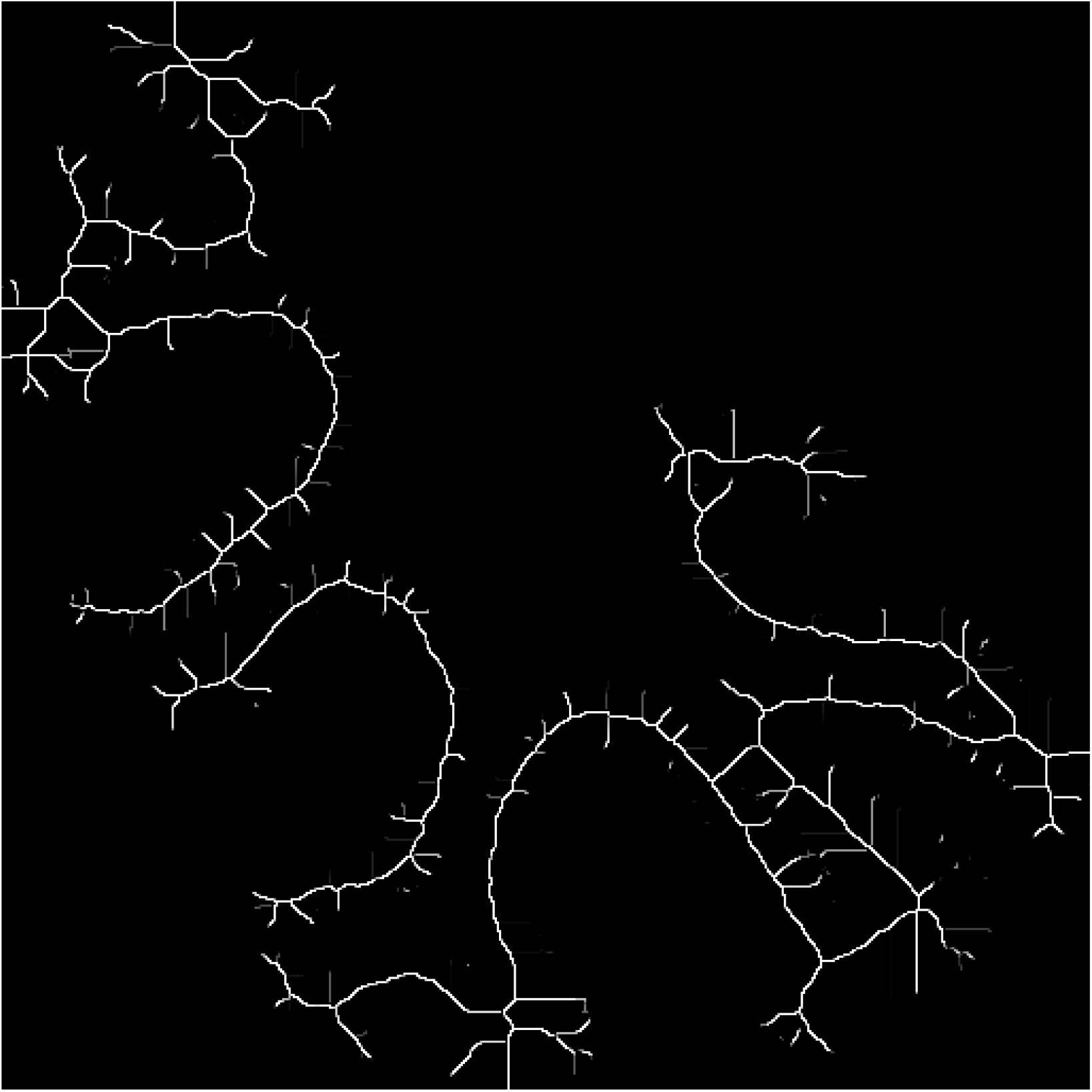
Morphological skeleton of the granule cell coordinates. The ST coordinates were scaled to a 512 × 512 binary image and a skeleton was estimated using the *magick* R package (Ooms, 2025). Although the skeletonization resembles the one-dimensional structure of the granule cells, the approach does not provide a way to convert this to a one-dimensional coordinate that can be used for downstream SVG detection.

**Figure S11.**
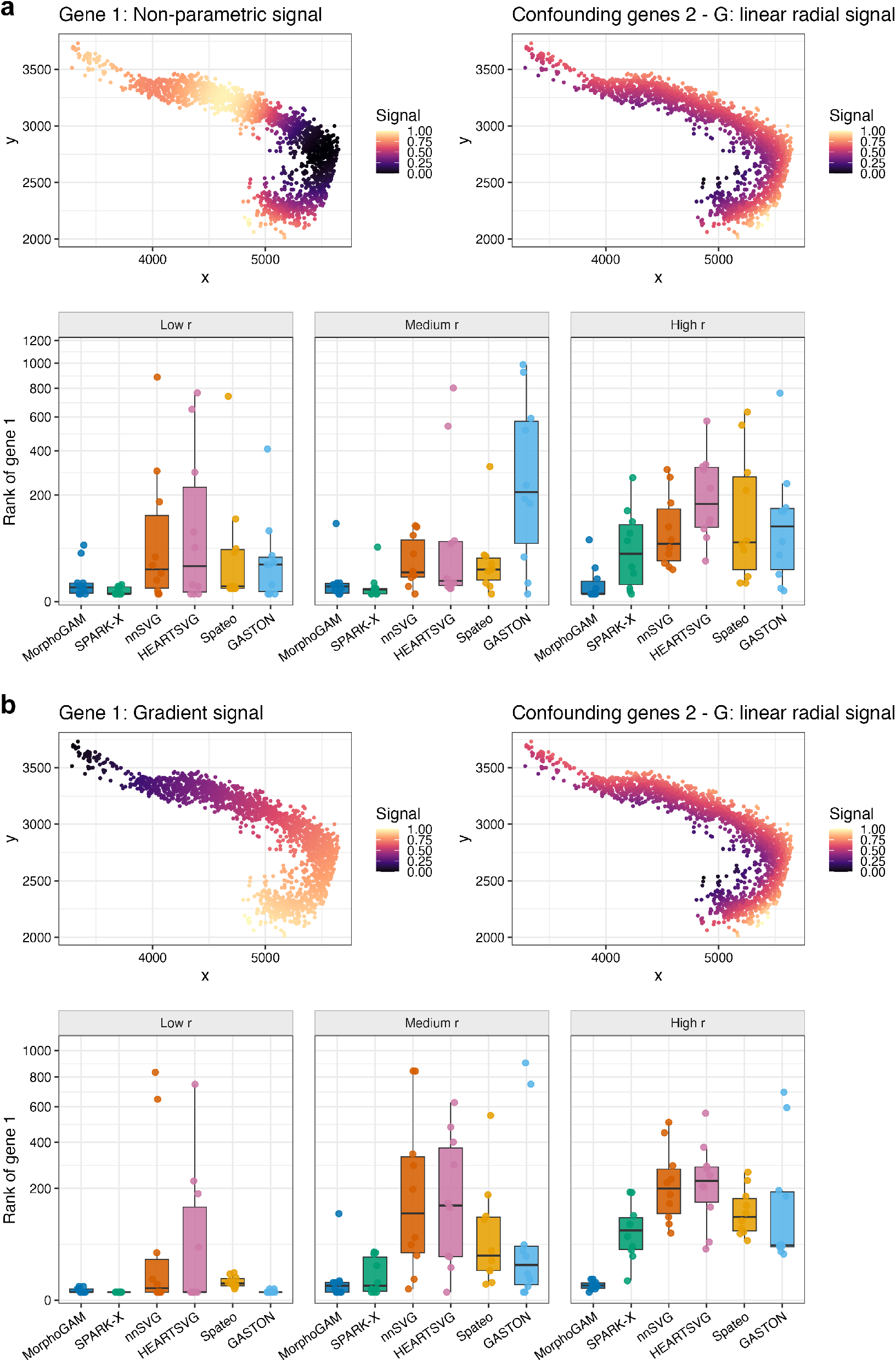
Repeating the simulation study of Figure 4d-e with different spatial patterns for the *t*-associated gene, see Methods for additional details.

**Figure S12.**
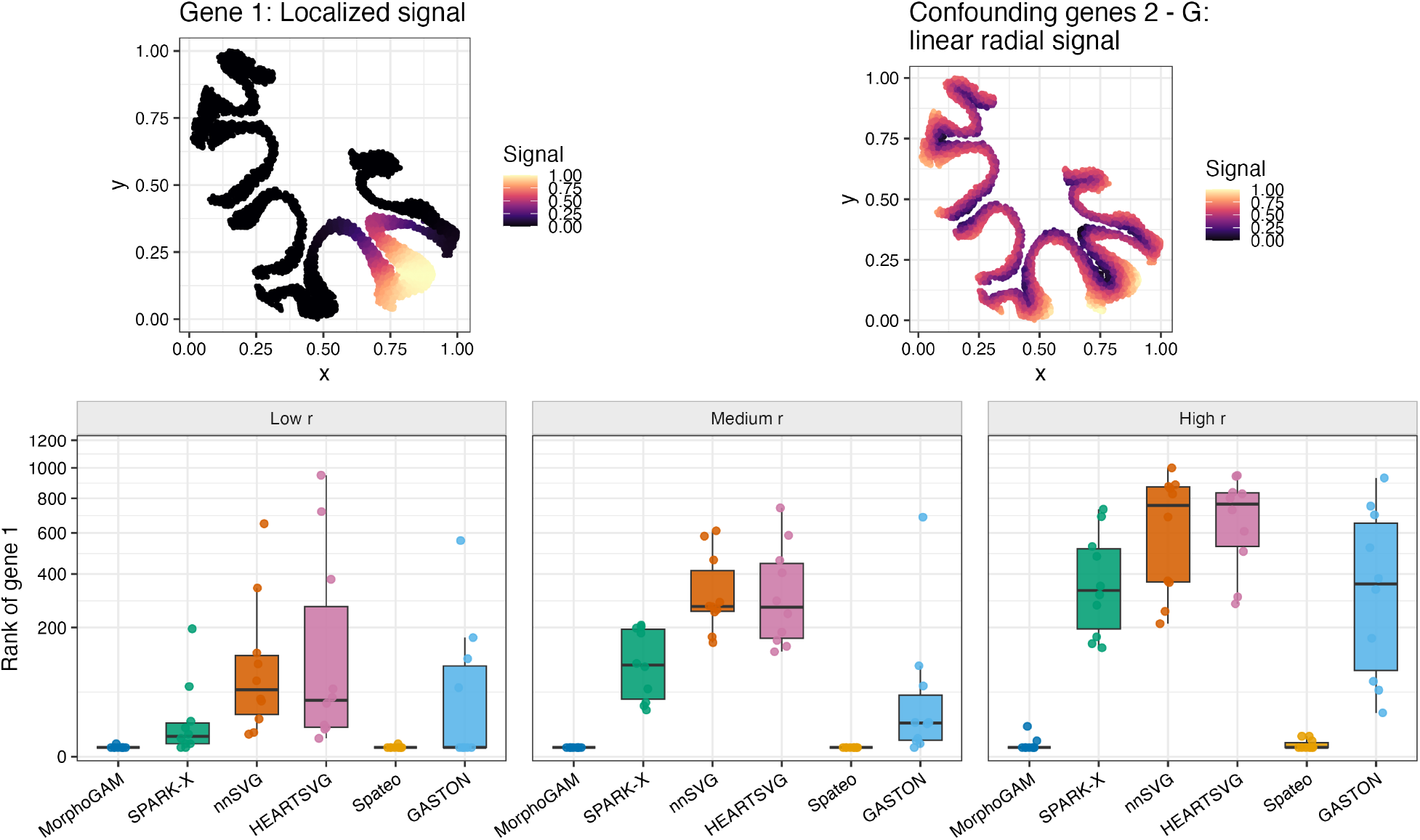
Repeating the simulation study of Figure 4d-e starting from the *x*-*y* coordinates of the granule cells.

**Figure S13.**
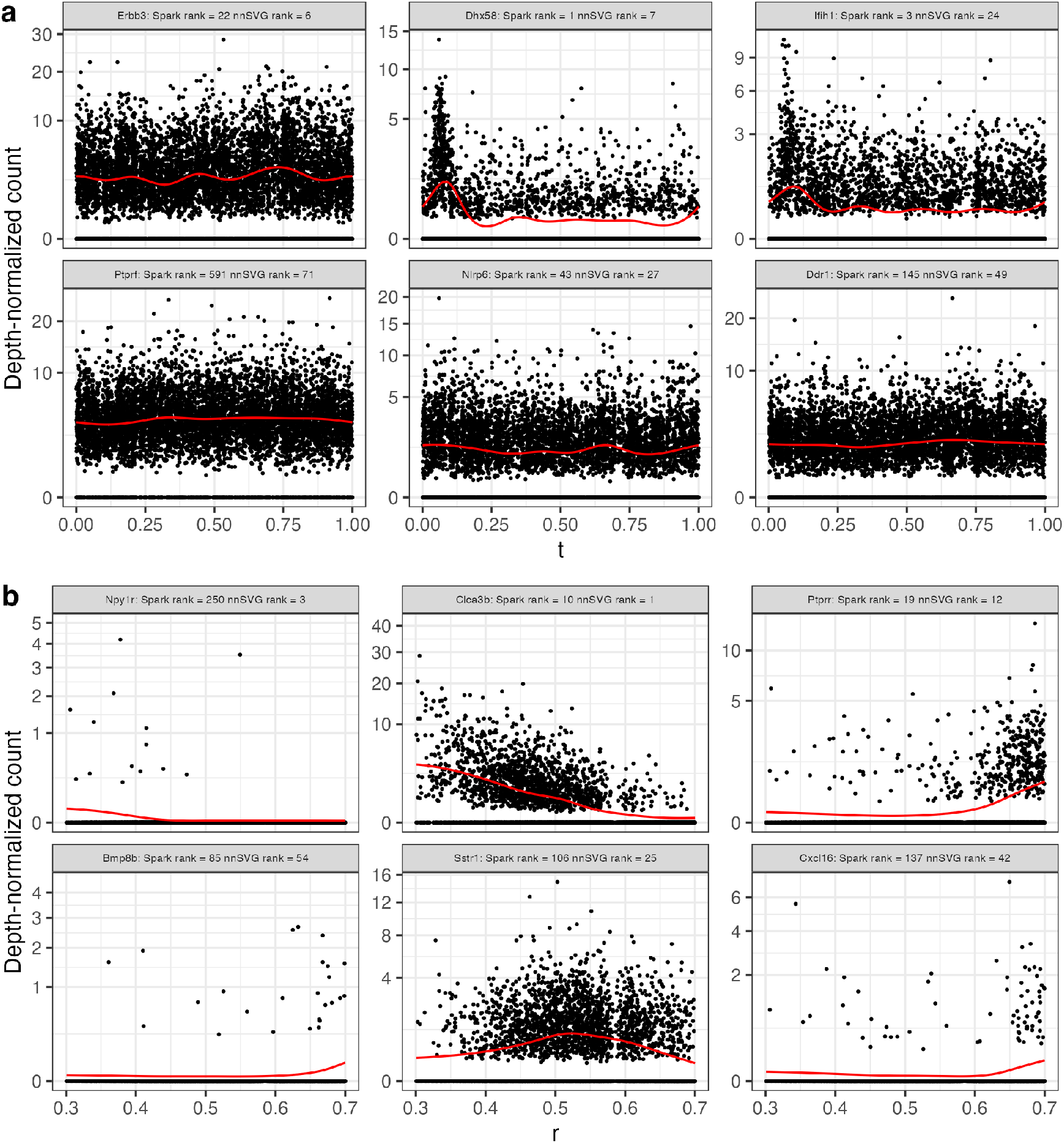
Repeating the analysis of Figure 5 plotting the genes with the largest range in the direction of the first morphologically relevant coordinate *t*_*j*_ and the the genes with the largest peak in the direction of the second morphologically relevant coordinate *r*_*j*_.

**Figure S14.**
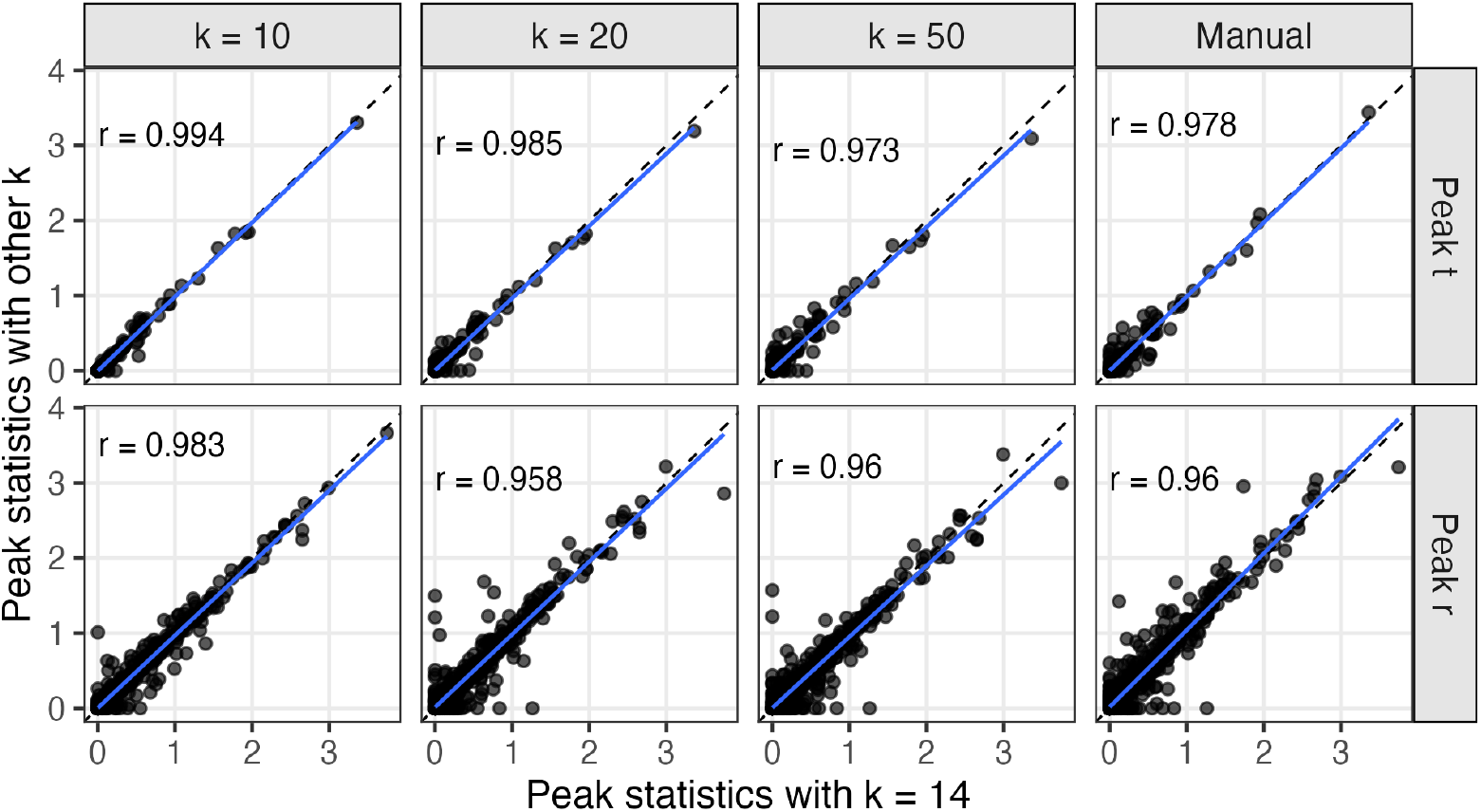
We repeated the SVG analysis in the mouse mucosa with different values of *k* used to estimate the curve *f* . We also included a manual (hand-drawn) curve. We computed the Pearson correlation between the peak statistics for each gene with *k* = 14 (the value used in the main text) with the peak statistics for the other choices of *k*.

**Figure S15.**
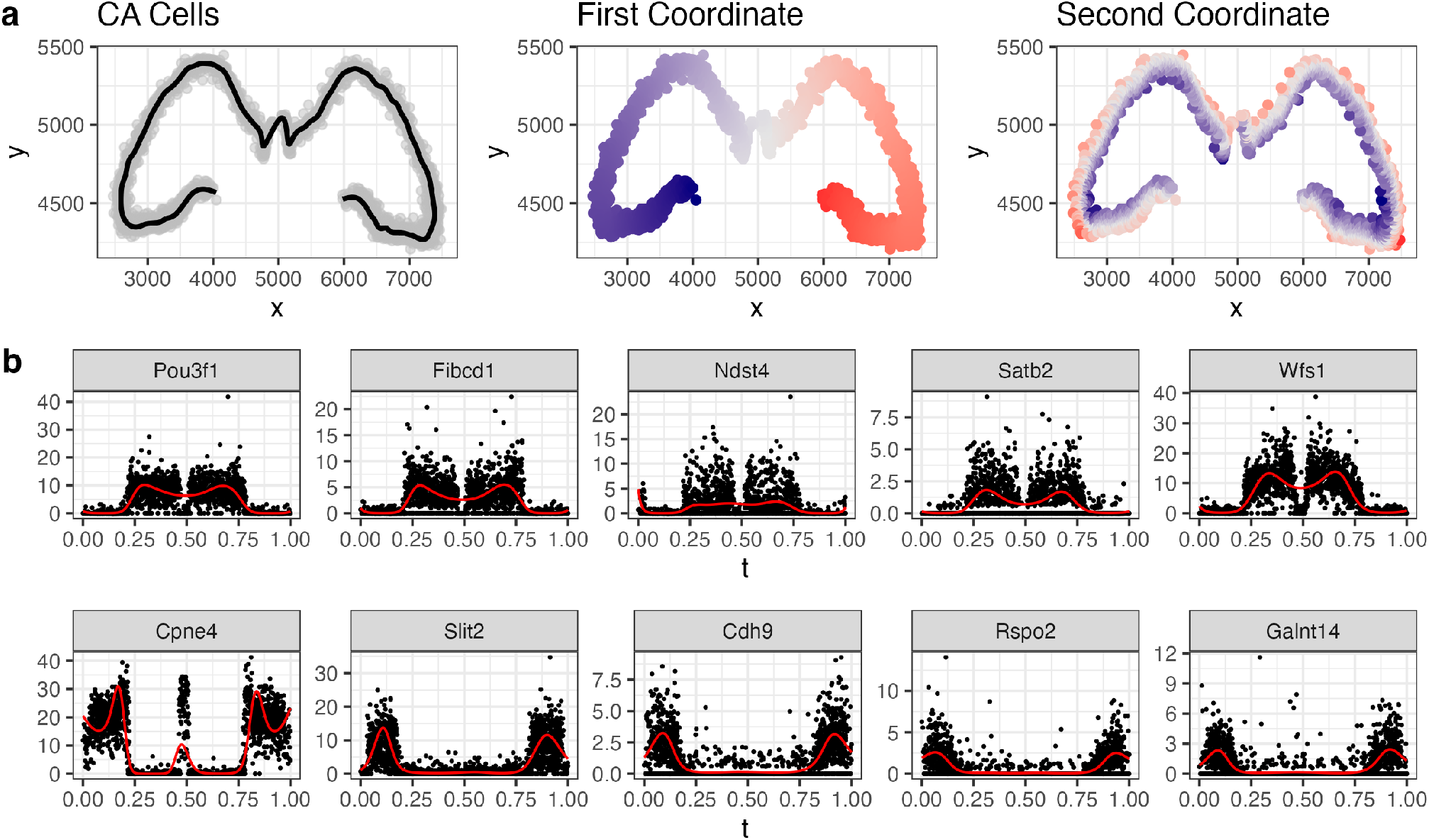
**a**. The spatial coordinates of CA1, CA2, and CA3 cells from a xenium dataset of the mouse brain. The first morphologically relevant coordinate traces the one-dimensional structure while the second morphologically relevant coordinate captures the orthogonal direction. **b**. The SVD-based approach (see Methods) was applied, and the genes with the largest *u*_*i*1_ (top row) and smallest *u*_*i*1_ (bottom row) were plotted. These genes are known markers of CA1 vs CA3 cells.

**Figure S16.**
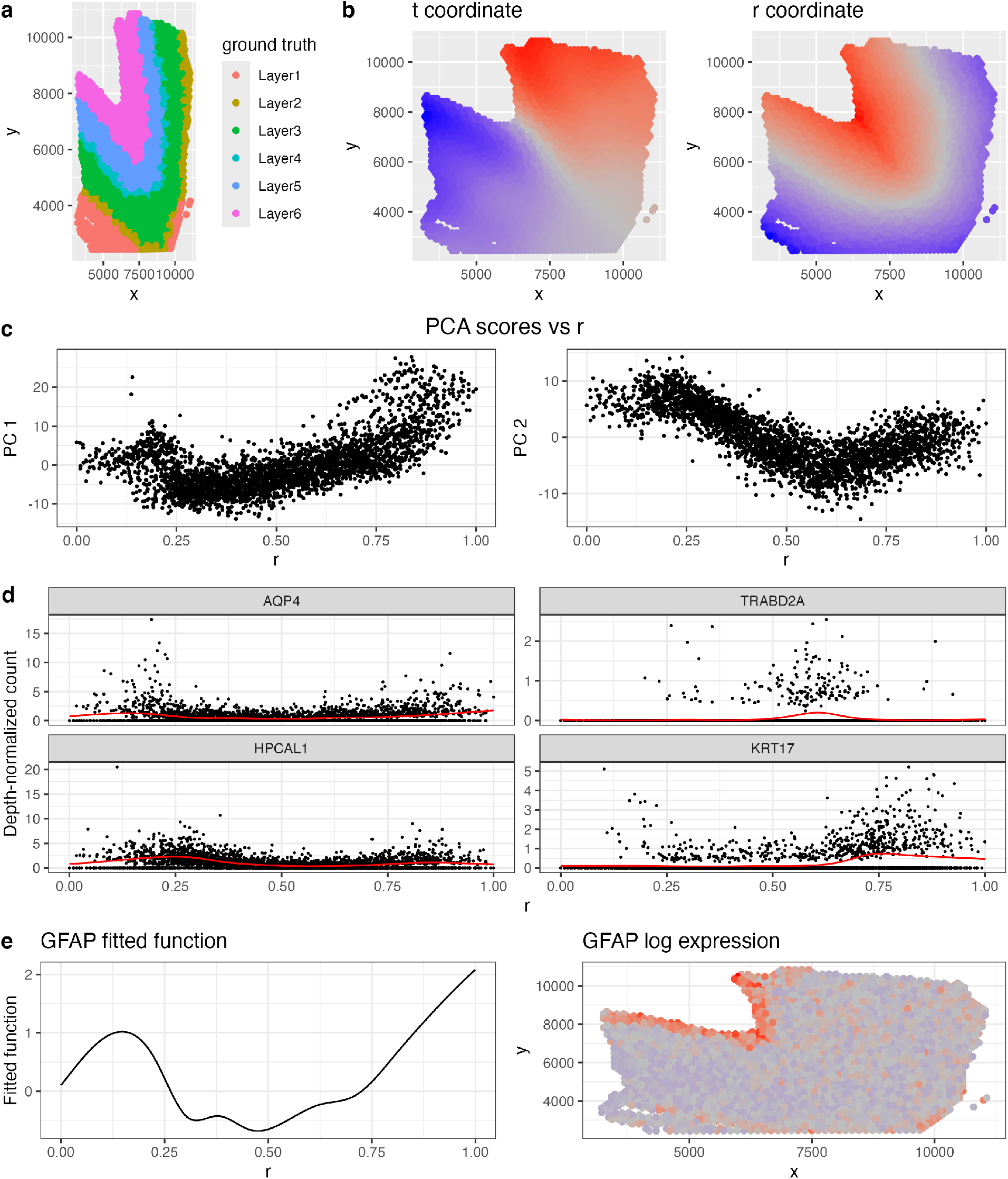
**a**. Spatial coordinates from a visium dataset of the dorsolateral prefrontal cortex (DLPFC) with cortical layers annotated by (Maynard et al., 2021). **b**. MorphoGAM was applied to identify a between (*t*) and across (*r*) layer coordinate system. **c**. We applied scGBM (Nicol and Miller, 2025), a count-based extension of PCA, to the subset genes with Bonferroni-adjusted *p*-value *<* 0.05 in the *r* direction. We then plotted the top two PC scores as a function of *r*, observing smooth variation. **d**. Novel layer-enriched genes identified by (Maynard et al., 2021) (Fig 4) were also identified to have variable expression when using the GAM model. **e**. MorphoGAM also identifies spatially variable expression of *GFAP* in a thin layer.

**Figure S17.**
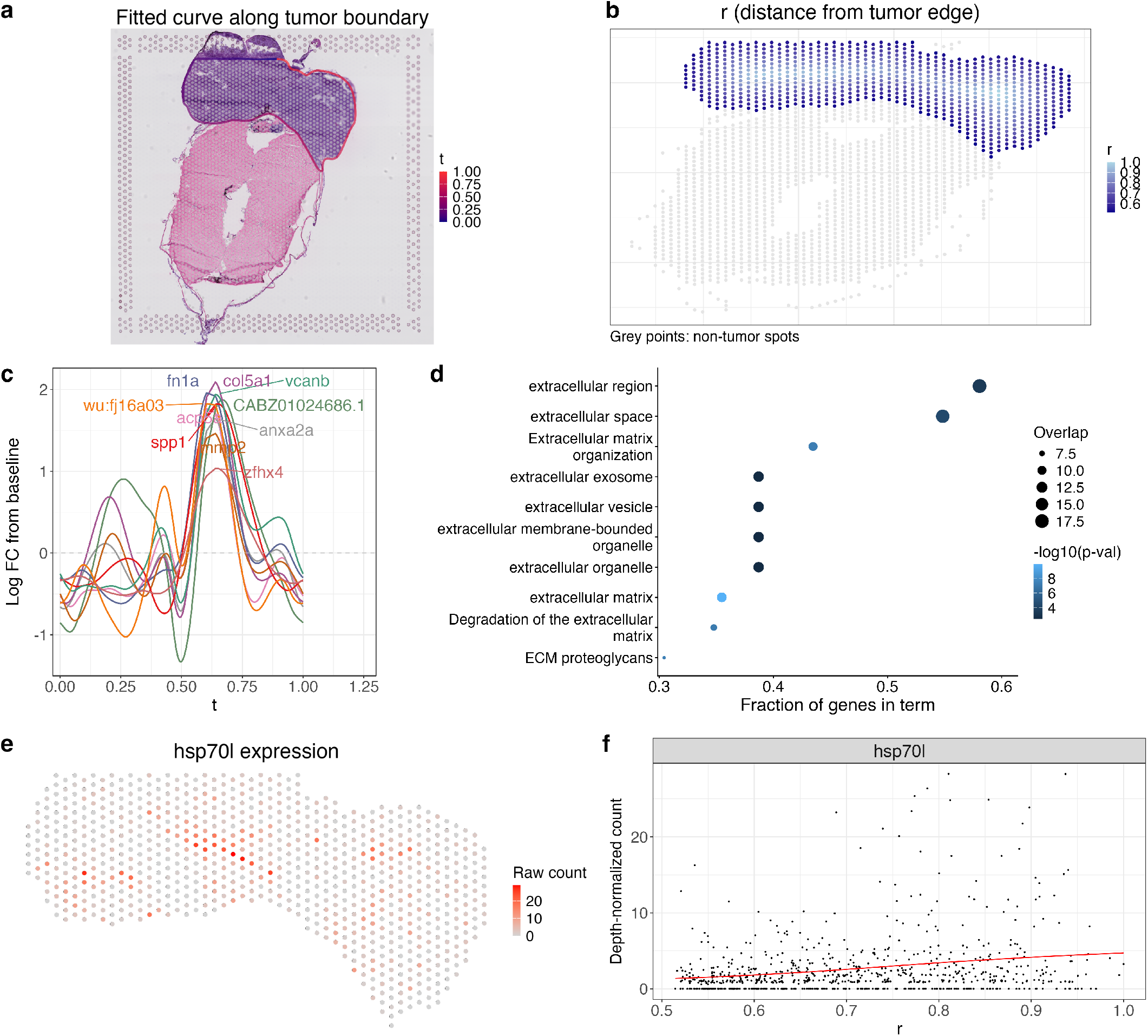
**a**. The fitted curve *f* (*t*) from MorphoGAM overlaid on the H&E image of Sample A from Hunter et al. (2021). **b**. *r* coordinate values are related to the distance from the spot to the tumor boundary. **c**. Applying SVD to the fitted *t* functions identifies a group of genes that have a peak near *t* ≈ 0.7. **d**. We applied gprofiler2 to the top 50 genes from the factor in the previous panel (Kolberg et al., 2020) to demonstrate over-representation of ECM-related genes. **e**. The raw counts for *hsp70l*. **f**. The fitted function in the *r*-direction for *hsp70l* (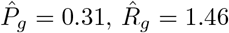, *p* = 0.00014).

**Figure S18.**
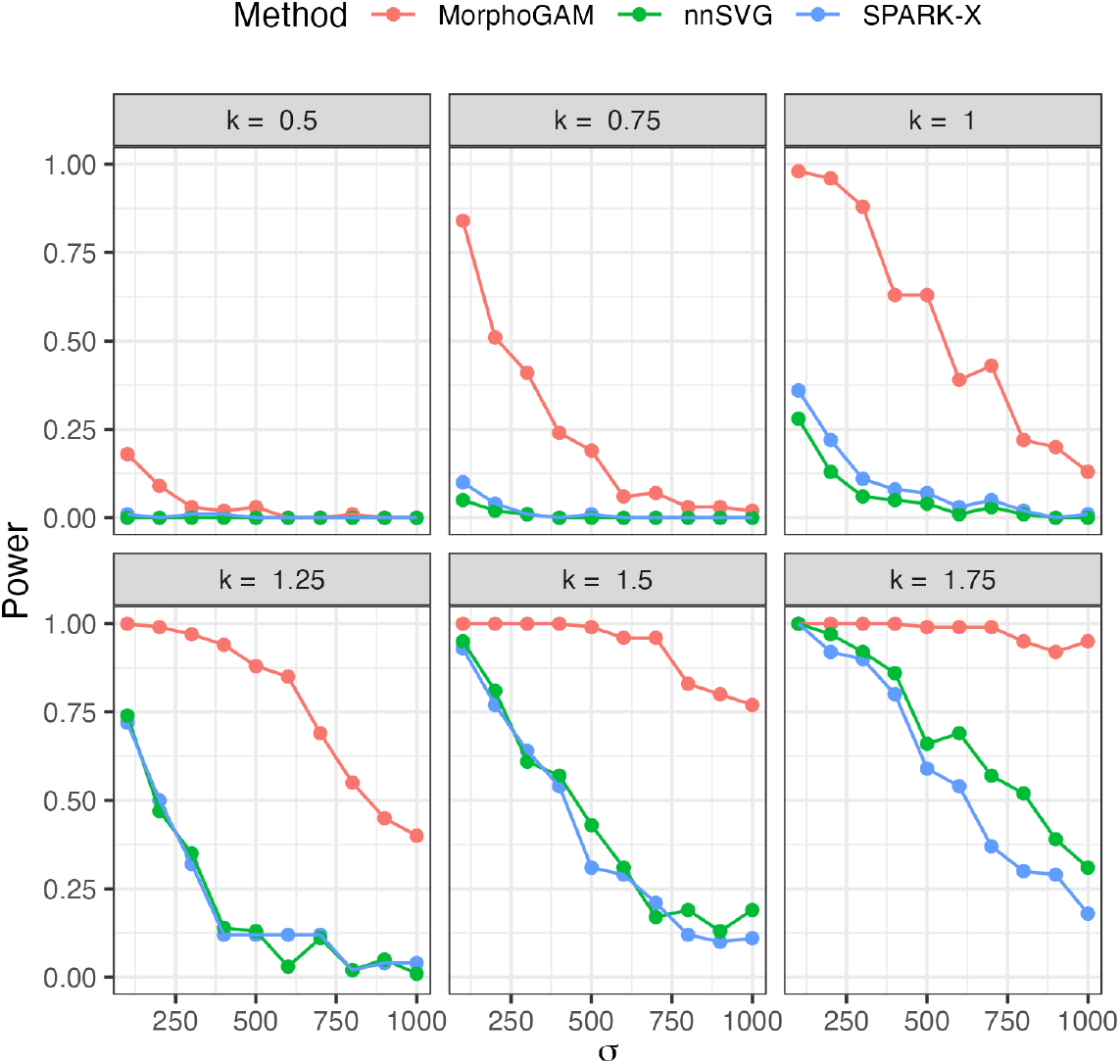
Comparing hypothesis-based frameworks to detect SVGs; a gene with 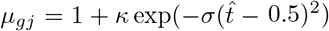 and *θ* = 5 was simulated and labeled as an SVG if the corresponding *p*-value was smaller than 0.05*/*20000. The power reflects the proportion of 100 trials where the null hypothesis was correctly rejected.

**Figure S19.**
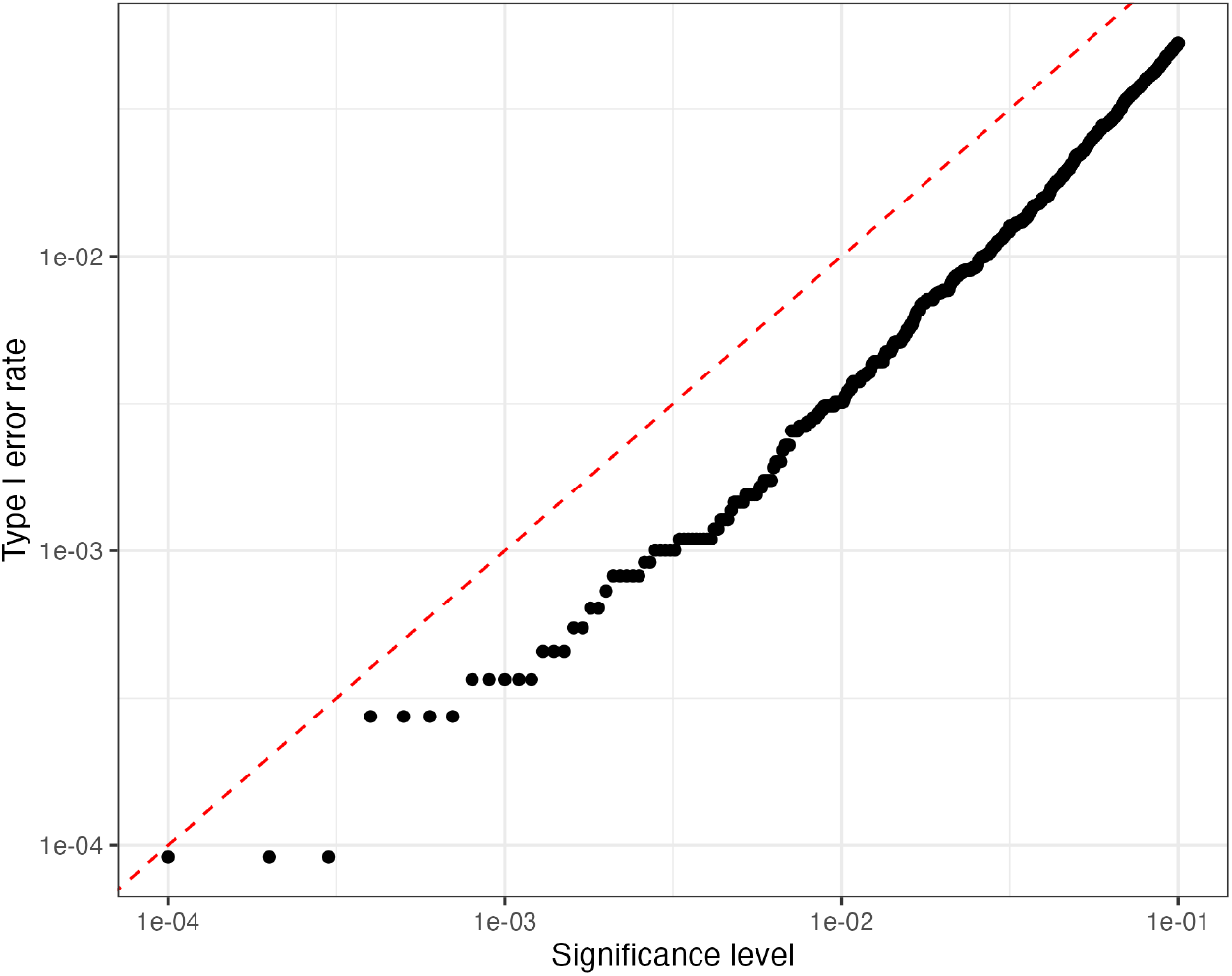
Spatial locations in the CA3 data were randomly permuted to produce a null dataset where there should be no SVGs. The proportion of genes with a *p*-value smaller than each significance level was computed (the Type I error rate). The red-dashed line indicates the nominal type I error rate.

**Figure S20.**
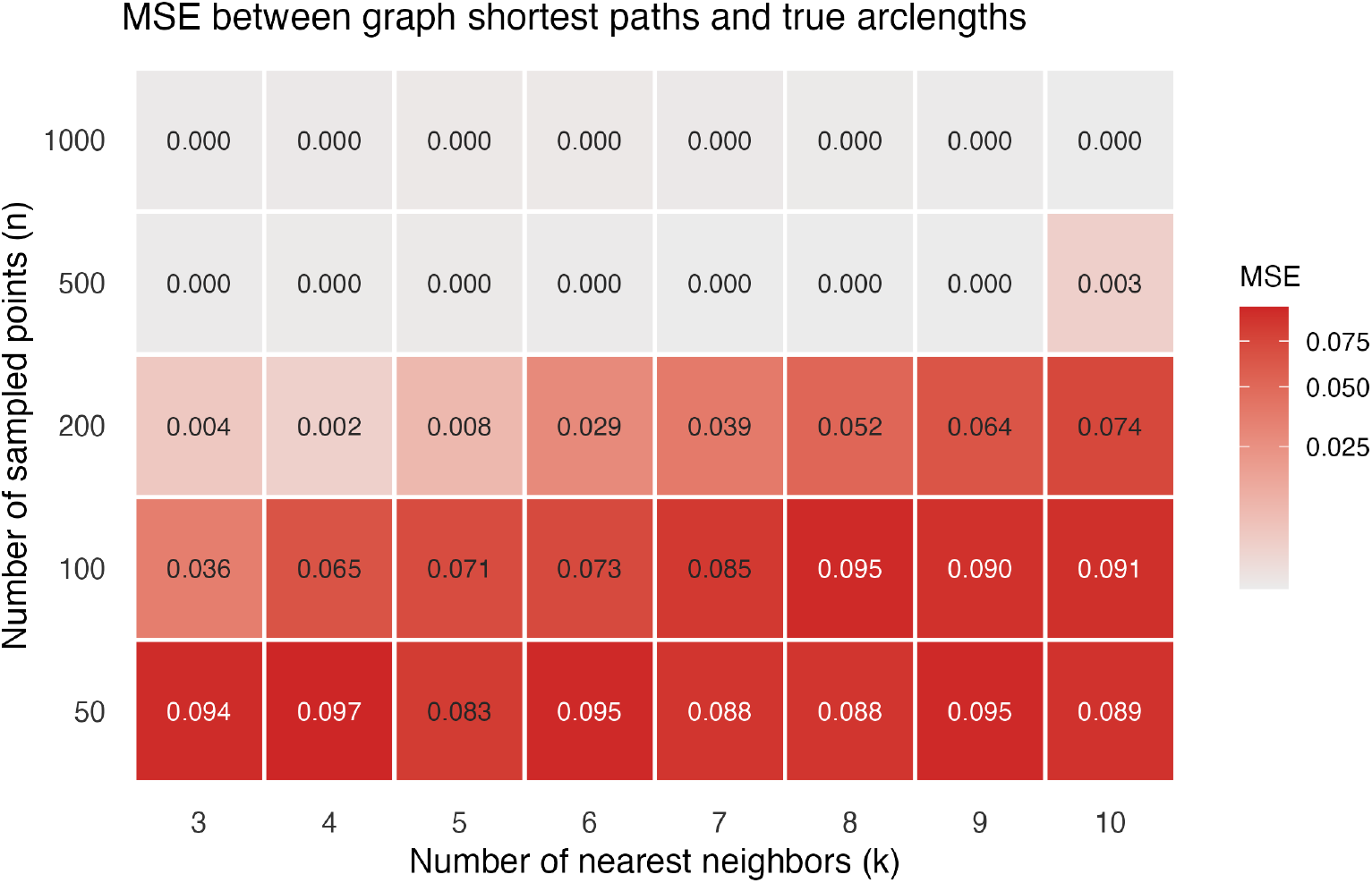
Using the swiss roll shape (see Figure S4), we computed the MSE between *k*-NN based shortest paths between two points to the true arclength, as the number of sampled points (*n*) and *k* are varied.

**Table S1.** Ratio diagnostic values by dataset. For the square, we drew 1000 points uniformly at random from the unit square [0, 1]^2^ (i.e., this is a structure that is not well approximated by a one-dimensional manifold).

| Dataset | Ratio Diagnostic |
| --- | --- |
| Colon (Figure 1) | 35.97 |
| Granule (Figure 3) | 34.26 |
| CA3 (Figure 4) | 8.69 |
| Square | 1.82 |

